# Systemic interindividual epigenetic variants link periconceptional environment to human fetal development

**DOI:** 10.64898/2026.06.12.731785

**Authors:** Chathura J. Gunasekara, Wen-Jou Chang, Maria S. Baker, Prachand Issarapu, Goo Jun, Garrett Hellenthal, Matt J. Silver, Andrew M. Prentice, Cristian Coarfa, Yumei Li, Rui Chen, Robert A. Waterland

## Abstract

At rare human genomic regions, DNA methylation states are established in the early embryo and maintained during cellular differentiation, yielding systemic (i.e. not tissue-specific) interindividual epigenetic variation. Previous screens for such correlated regions of systemic interindividual variation (CoRSIVs) were limited to White Americans. Here, we describe the first human CoRSIV screen including self-identified Black and White Americans. We integrate deep whole-genome bisulfite sequencing data for three tissues from each of ten Black and ten White donors in the NIH Genotype-Tissue Expression program. This approach identifies twice as many CoRSIVs among Black than White Americans. Establishment of CoRSIV methylation is sensitive to periconceptional environmental exposures including assisted reproduction, seasonal variation, and famine. CoRSIV-associated genes are enriched for GWAS variants linked to cancer and neurodevelopment. Although only 15% of Black CoRSIVs overlap with those among White individuals, both sets are associated with the same subfamilies of transposable elements. Within multiple cell lines, ranked enrichments of transcription factor binding to Black and White CoRSIVs are exquisitely coordinated and related to genome organization, indicating that CoRSIV methylation states established in the early embryo play an important role in guiding subsequent cellular differentiation.

## Main text

Methylation of CpG dinucleotides in DNA, the most stable epigenetic mark in mammals, plays essential roles in controlling parent-of-origin-specific expression of imprinted genes, silencing transposable elements, and orchestrating cellular differentiation. Most interindividual variation in DNA methylation is limited to specific tissues or cell types, complicating population-level association studies of epigenetic variation and risk of disease. At correlated regions of systemic interindividual epigenetic variation (CoRSIVs), however, interindividual epigenetic variation occurs systemically, i.e. consistently across diverse organs and cell types^1^. Systemic interindividual variation (SIV) occurs when DNA methylation states are established in the cleavage-stage embryo then propagated during subsequent rounds of cell division, organogenesis, and cellular differentiation. Early embryonic establishment of CoRSIV methylation is influenced by both genetic variation^2^ and the periconceptional environment^1,3–5^, implicating CoRSIV^TM^ methylation as a potential mediator of developmental origins of disease^6^. CoRSIVs are advantageous for epigenetic epidemiology because measuring their methylation in easily obtainable tissues like peripheral blood provides information about epigenetic regulation throughout the body^7,8^. Additionally, owing to their early-embryonic establishment and long-term stability, CoRSIV methylation states measured in blood may be lagging indicators of epigenetic states that were established prior to and potentially influenced organogenesis and differentiation of every organ and cell type in the body.

Whereas SIV is associated with sub-telomeric regions, specific subfamilies of transposable elements, and single-nucleotide variants (SNVs)^1,2^, its precise genomic determinants remain poorly understood. Beyond influencing the extent of methylation at associated CoRSIVs, genetic variation amongst diverse population groups could also determine whether a specific genomic region is a CoRSIV. However, whilst many CoRSIVs identified in White Americans also exhibit SIV in Asian^4^ and African^3^ individuals, all previous human screens for SIV^1,4,9^ were based on White individuals in the US, so there is a great need to determine the extent to which SIV differs across different self-identified racial groups. Here, we addressed this need by expanding our previous CoRSIV screen to include self-identified Black Americans, advancing our understanding of interindividual epigenetic variation in individuals of different ancestry.

## Results

### Twice as many CoRSIVs among Black than White Americans

We previously developed an unbiased genome-wide approach to identify CoRSIVs^1^ using deep whole-genome bisulfite sequencing (WGBS) to profile CpG site-specific DNA methylation in three tissues in each of 10 self-identified White GTEx^10^ donors (CoRSIVs 1.0). We’ve now expanded the screen to include WGBS data on brain, heart, and thyroid (ectodermal, mesodermal, and endodermal lineages, respectively) from each of 10 self-identified Black or African-American GTEx donors (hereafter referred to as Black) (Fig. 1a; Supplementary Table S1). The ∼40x per tissue average sequencing depth of the WGBS data for each of the Black donors (Supplementary Table S2) is comparable to that generated for the White donors^1^. We also employed the same analytical pipeline^1^, with one important refinement. Because deamination of methylated cytosines yields thymine, and CpG sites are the predominant site of methylation in the human genome, C to T transitions at CpG sites (and corresponding G to A transitions on the opposite strand) are the most common SNV in mammalian genomes^11^. In bisulfite-converted, whole-genome amplified DNA, unmethylated cytosines are indistinguishable from C to T transitions. Our original screen^1^ did not consider this. We did evaluate this issue in our subsequent validation studies using target-capture bisulfite sequencing^2^, but due to an analytical error underestimated the extent of CpG SNVs within the targeted regions. Here, revisiting our 2023 analysis of 3,914 genic CoRSIVs and analyzing GTEx genotyping data on the 188 donors showed that most CoRSIVs 1.0 contain two or more SNVs (Supplementary Fig. 1a). Moreover, most of these affect CpG sites; in over half of CoRSIVs 1.0 two or more CpGs are affected by a SNV (Supplementary Fig. 1b). In many cases, these CpG-SNVs drove much of the variation that was interpreted as CoRSIV-average methylation (two examples are shown in Supplementary Fig. 1c). Therefore, here we included an additional step to eliminate this contaminating genetic variation. We used data from the 1000 Genomes Project to identify common SNVs overlapping CpG sites among White (CEU) and Black (ASW) Americans. Then, utilizing genotype calls from GTEx, we identified all such CpG sites at which one or more of the ten individuals in either screen carries a variant allele. All such variant CpG sites were masked in all individuals for the remainder of our computational approach.

**Fig. 1.**
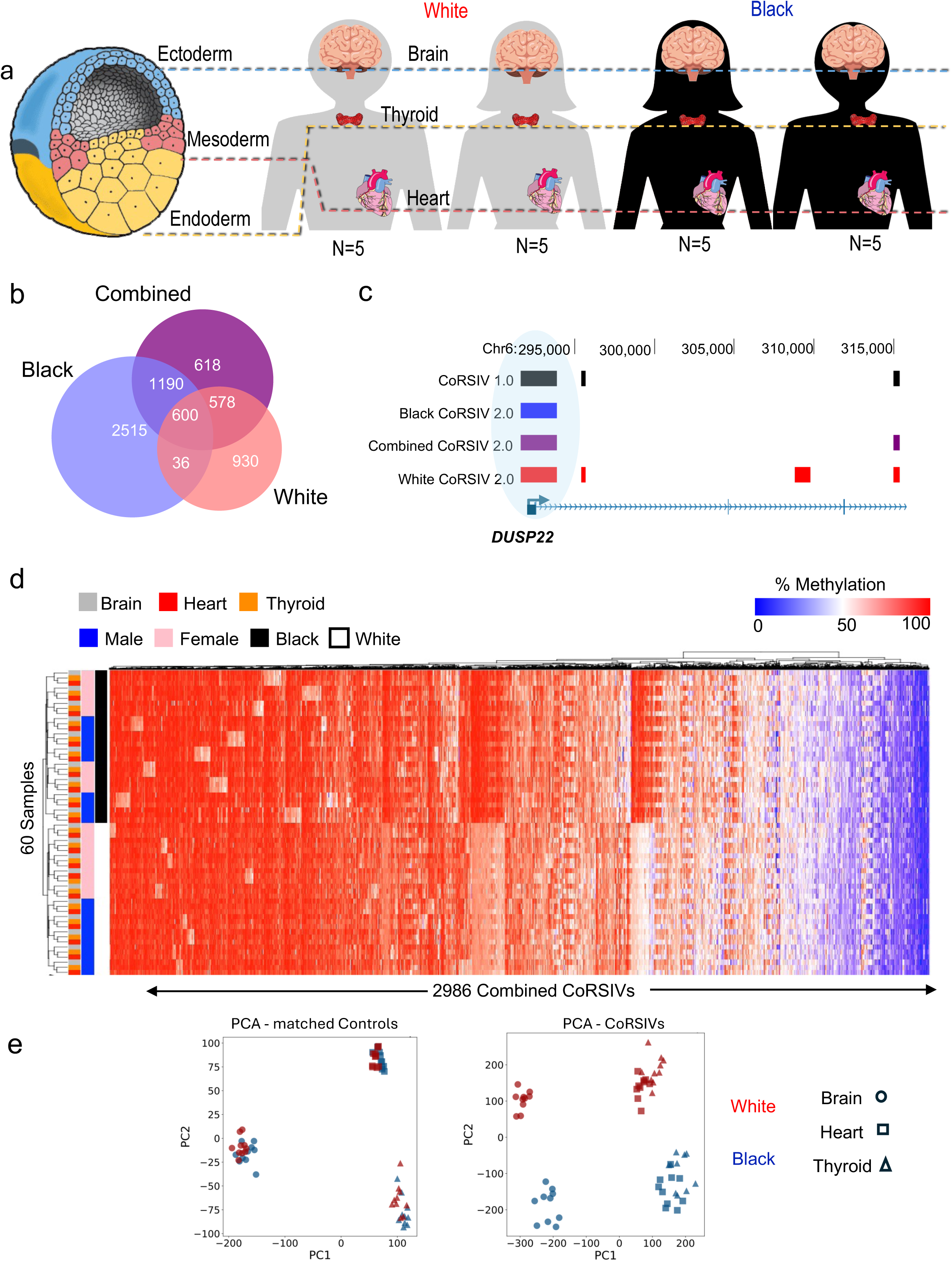
A multi-racial screen for systemic interindividual epigenetic variation (SIV). **a** The tissues analyzed, brain, thyroid, and heart, represent the three embryonic germ layer lineages; 10 White and 10 Black GTEx donors were profiled by whole-genome bisulfite sequencing, yielding 60 methylomes. **b** SIV was assessed across all 20 individuals (Combined) and amongst the 10 Black or 10 White individuals only, yielding overlapping subsets of CoRSIVs. Only 15% of CoRSIVs identified among Black individuals overlap with those identified among Whites. **c** Those that overlap, however, match almost perfectly, as at exon 1 of *DUSP22*. Note also the CoRSIV in *DUSP22* intron 3, detected amongst both CoRSIVs 1.0 and 2.0. **d** Unsupervised clustering of average methylation data for the 2,986 combined CoRSIVs groups libraries by individual as well as by race. **e** Principal component analysis (PCA) of methylation at matched control regions (left) clusters samples by tissue, but not by race. PCA of methylation at CoRSIVs (right) clusters samples by race and additionally separates brain from heart and thyroid.

Focusing on all 100 bp genomic bins containing one or more CpG sites, we employed a two-step analytical approach to maximize genomic coverage. First, combining reads across the three tissues from each individual, we calculated bin-level individual average methylation values and constructed blocks of individually-correlated methylation based on correlations amongst adjacent bins. Then, using the tissue-specific data, we tested each such block for systemic interindividual variation (SIV) by calculating inter-tissue correlations. To capture all variation in the sample we conducted three separate screens, one each on the ten Black donors, the ten White donors, and the 20 donors combined, detecting a total of 39,368 regions. Using previously established cutoffs to target the most biologically meaningful variants^1^, we focused subsequent analyses on the 6,467 unique regions each containing ≥5 CpGs and exhibiting an interindividual average methylation range ≥20% (Supplementary Fig. 2a). This approach identified 4,341, 2,144, and 2,986 CoRSIVs in the Black, White, and combined screens, respectively (Fig. 1b) (together, hereafter referred to as CoRSIVs or CoRSIVs 2.0). Of White CoRSIVs 2.0, 83% overlap with CoRSIVs 1.0 (Supplementary Fig. 3). WGBS summary statistics, genomic coordinates (hg38) and gene annotations of CoRSIVs 2.0 are provided in Supplementary Table S3.

Despite the two screens being equally powered, twice as many CoRSIVs were identified among Black than White donors. Further, only 636 were identified by both screens (Fig. 1b). This low overlap is not attributable to inadequate power. Underscoring the reliability of the WGBS data, most (63%) CoRSIVs identified in both Black and White groups showed complete overlap (Supplementary Fig. 2b) including, for example, the 2,200bp CoRSIV encompassing *DUSP22* exon 1 (Fig. 1c). Though the combined screen included the most individuals and the greatest genetic variation, it captured fewer CoRSIVs than the screen limited to the 10 Black donors; moreover, 80% of them were also detected in one or both of the race-specific screens (Fig. 1b). To better understand the unique genomic characteristics of CoRSIVs, we generated two set of control regions for comparison^1^. To represent genomic background, 10,000 random control regions were drawn randomly from across the genome, matching the genomic size (bp) distribution of the 6,467 CoRSIVs 2.0 (Supplementary Table S4). Then, to evaluate the extent to which genomic associations of CoRSIVs are driven by their CpG density, we generated a set of matched control regions^1^; for each of the 6,467 CoRSIVs 2.0 we selected a control region matched on chromosome, region size, and number of CpGs (Supplementary Table S5). To enable race-specific comparisons, we also generated sets of random and matched controls based on only the Black or White CoRSIV sets.

As for CoRSIVs 1.0^1^, unsupervised hierarchical clustering of average methylation across the 2,986 combined CoRSIVs (Fig. 1d) grouped the 60 WGBS libraries perfectly by individual. They also grouped by self-identified race; two clusters of CoRSIVs with obvious racial differences show higher methylation in the Black vs. White donors, consistent with previous array-based analyses of DNA methylation in blood^12^. At matched control regions, by comparison (Supplementary Fig. 4), the 60 WGBS libraries grouped by tissue and, within tissue, nearly perfectly by race. In a principal component analysis (PCA) of methylation in the matched control regions (Fig. 1e, left) the effect of tissue is dominant, with little separation by race. The PCA for CoRSIVs identified in the Combined screen, however, distinguishes samples by race (Fig. 1e, right). Within each race, heart and thyroid cluster together but, as previously observed^2^ the brain (cerebellum) samples are distinct, despite high inter-tissue correlations. These results indicate that compared to control regions with similar CpG density, CoRSIVs more effectively capture racial differences in DNA methylation.

### Genomic Characteristics of Black and White CoRSIVs

The genomic distribution of CoRSIVs 1.0 is non-uniform^1^, exhibiting pronounced enrichment at the MHC locus on chromosome six and a pericentromeric region on chromosome 20. CoRSIVs 2.0 show similar enrichments, in both races (Fig. 2a). One obvious racial difference, however, is a region of high CoRSIV density at the chromosome 9 pericentromeric region specifically among Black individuals. Similar analysis of matched control regions (Supplementary Fig. 5a) shows they too are enriched at the MHC locus, indicating that its high CoRSIV density may be attributable to its CpG density. CpG density does not, however, appear to explain the enrichment of CoRSIVs in the pericentromeric region of chromosome 20. Black matched controls do show a show high density in the pericentromeric region of chromosome 9 (Supplementary Fig. 5b), corresponding to the peak of Black CoRSIV density. Random control regions matched to Black or White CoRSIVs, however, show no obvious differences (Supplementary Fig. 5c). To understand why some genomic regions qualify as CoRSIVs in one race but not the other, we plotted out the WGBS data in a subset of such regions. One on chromosome 8 encompasses the first exon of a long non-coding RNA and is a CoRSIV (shaded area) among Black but not White individuals (Fig. 2b). This region shows marked systemic interindividual variation involving several of the Black donors but little variation among White individuals. In most cases, when a CoRSIV is detected among Black but not White donors, there is some interindividual variation among the White donors but it is both less pronounced and not systemic (i.e. inconsistent across the three tissues) (Supplementary Fig. 6a,b). In other cases (Supplementary Fig. 6c,d), CoRSIVs exhibit interindividual variation in Black individuals but little or no variation in White individuals.

**Fig. 2.**
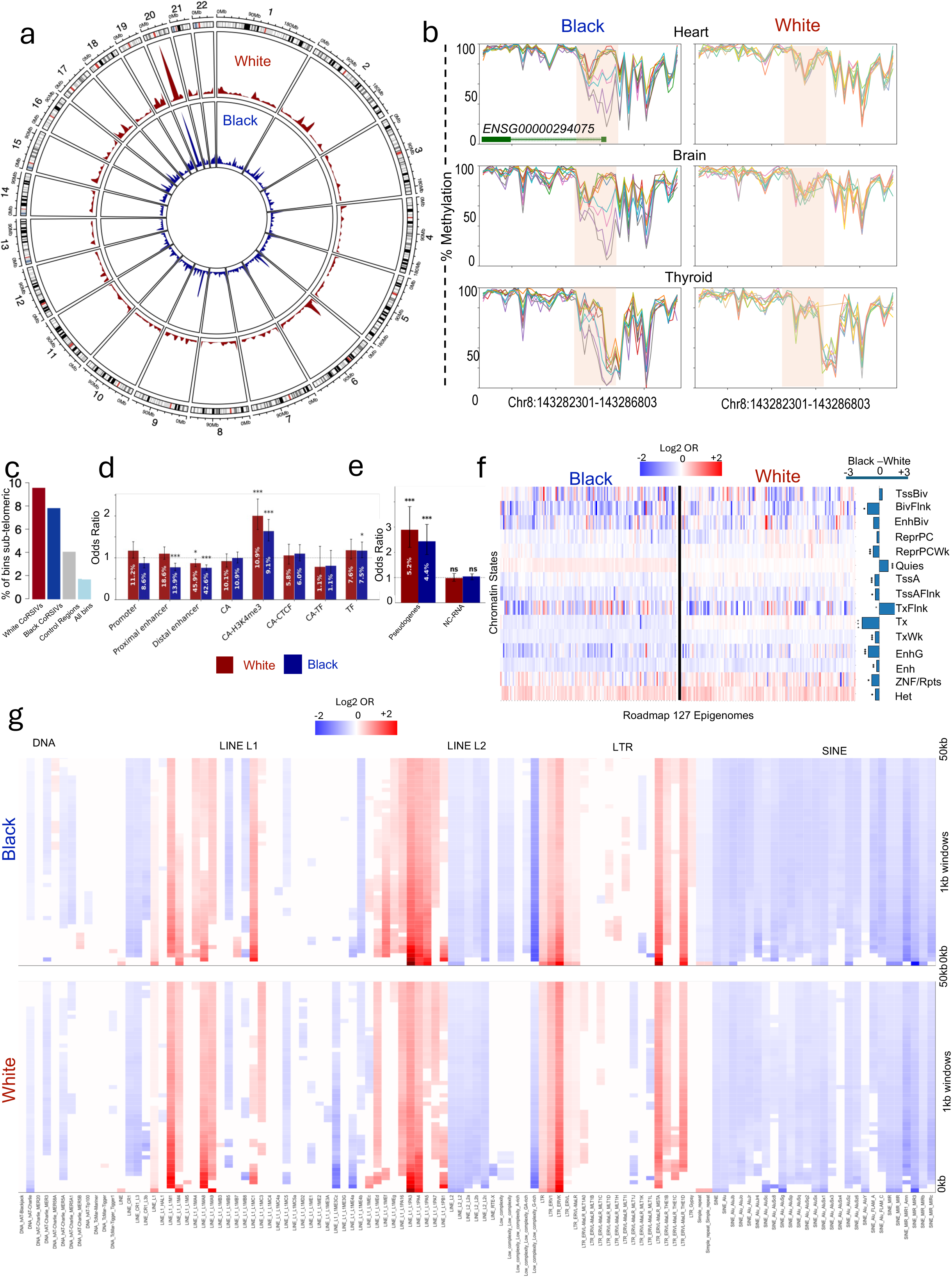
Distribution and genomic features of CoRSIVs in Black and White Americans. **a** circos plot of human autosomes depicts CoRSIV density among White and Black Americans. **b** Tissue-specific, individual-level methylation data for a chr8 CoRSIV detected among Black but not White individuals. The CoRSIV (shaded area) overlaps the 5’ end of the lncRNA *ENSG00000294075*. **c** Compared to all informative bins, White and Black CoRSIVs are ∼five-fold and ∼four-fold enriched in sub-telomeric regions, respectively (Chi-sq P < 10^-300^). **d** Analysis of ENCODE candidate cis-regulatory elements. Relative to matched control regions, Black and White CoRSIVs are ∼1.8-fold enriched for overlap with H3K4me3 in accessible chromatin (CA-H3K4me3) (Chi-sq P = 5.7x10^-10^, 2.3x10^-13^) . **e** Compared to matched control regions, neither Black nor White CoRSIVs are enriched for ncRNAs. But, both Black (OR:2.4; P=3.4x10^-13^) and White (OR:2.9; P=2.7x10^-13^) CoRSIVs are enriched for overlap with pseudogenes. **f** Analysis of CoRSIV colocalization with 15 ChromHMM chromatin states (rows) learned from 127 Roadmap Epigenomes (columns). Most group differences (Black - White) are minor, as summarized by the bar plots (right). Black but not White CoRSIVs are 52% enriched within the quiescent state (Quies) comprising 68% of the genome. **g** Black (top) and White (bottom) CoRSIV-flanking regions show long-range enrichments and depletions for specific families and subfamilies of transposable elements. Using 1 kb step sizes (rows), colored cells in each column indicate statistically significant enrichments or depletions for subfamilies within each of 7 classes of transposable element within +/- 50 kb of CoRSIVs. Genomic regions flanking CoRSIVs identified among Black and White Americans show similar long-range depletion of SINE and LINE-2 elements and both enrichments and depletions in specific subfamilies of LINE-1 and LTR elements.

Despite their modest overlap, genomic characteristics of Black and White CoRSIVs 2.0 are similar. For example, as in our original report^1^, CoRSIVs 2.0 are enriched in subtelomeric regions (Fig. 2c), relative to either random or matched control regions (Chi-sq test p < 10^-300^ for each comparison). Analyzing ENCODE candidate cis-regulatory elements^13^ we found that, compared to matched control regions, CoRSIVs in Black and White Americans are 1.5 to 2-fold enriched for H3K4me3 marks in accessible chromatin (Fig. 2d) (P= 5.7x10^-10^ and P=2.0x10^-13^, respectively). Summary statistics of the candidate cis regulatory element analysis are provided (Supplementary Table S6). Observing that many CoRSIV 2.0 regions are near pseudogenes and non-coding RNA (ncRNA), we evaluated associations with these features. Compared to matched control regions, there was no enrichment for ncRNA, but both White and Black CoRSIVs showed a two to three-fold enrichment for pseudogenes (Fig. 2e; Supplementary Table S7) (P= 2.7x10^-13^ and P= 3.45x10^-13^, respectively). Evaluating CoRSIVs in the context of ChromHMM chromatin states identified by analysis of Roadmap Epigenomics Consortium data^14^ showed generally similar associations with Black and White CoRSIVs; both are enriched for heterochromatin (Het) and depleted for enhancers (Enh). Minor racial differences were observed, however. Most notably, Black but not white CoRSIVs are depleted in regions of strong transcription (Tx) and enriched in quiescent states (Quiesc) that comprise a substantial proportion of the genome (Fig. 2f). The log_2_ odds ratios shown in the heatmap are provided (Supplementary Table S8). Using the Gencode 48 database, we calculated the percentage of CoRSIVs within +/- 3KB of transcription start sites (TSS) and transcription end sites (TES) and overlapping gene bodies (Supplementary Fig. 7). Relative to matched control regions, both Black and White CoRSIVs are over-represented in intergenic regions and under-represented in gene bodies (Chi-sq test p<0.001). Regardless, nearly 70% of either Black or White CoRSIVs are gene-associated (i.e. overlapping or within 3kb of a gene).

Most strikingly, although only 15% of CoRSIVs identified in Black Americans overlap those identified in Whites (Fig. 1b), patterns of enrichment and depletion of transposable elements in CoRSIV-flanking regions are highly similar across the two sets of regions (Fig. 2g). Consistent with our previous report^2^, compared to matched control regions, the 50kb regions upstream and downstream of both Black and White CoRSIVs 2.0 are depleted for LINE2 and SINE elements and enriched for specific sub-families of LINE1 elements and long-terminal repeats (LTRs) (Fig. 2g). In particular, genomic regions flanking CoRSIVs 2.0 are enriched for L1M1, L1M4, L1MA8, L1MA9, L1PA3, and L1PA4 elements and, in the LTR family, ERVK, MalR-MSTA and MalR-THE1D (Fig. 2g; Supplementary Table S9). Long-range enrichments for these same sub-families were observed for CoRSIVs 1.0^2^. Overall these analyses show that, although CoRSIVs identified in Black and White Americans are largely distinct, they generally associate with the same genomic features.

### Assessment of genetic influences on CoRSIVs 2.0

We previously used target capture bisulfite sequencing to profile DNA methylation at over 4,000 gene-associated CoRSIVs in multiple tissues from up to 188 GTEx donors^2^ (Fig. 3a,b). Here, we re-analyzed these data focusing on the 774 CoRSIVs 2.0 with adequate sequencing depth. High inter-tissue correlation is the hallmark of systemic interindividual variation^15^. For example, the target capture data on one CoRSIV (Fig. 3c) illustrates that methylation in five different tissues is strongly correlated with that in blood. A heat map summarizing all possible inter-tissue correlations across the 774 informative CoRSIVs (Fig. 3d) shows generally strong positive inter-tissue correlations, validating SIV at these regions. (For summary statistics see Supplementary Table S10). Availability of GTEx SNV genotyping data for the profiled subjects allowed us to assess genetic influences on CoRSIV methylation (cis methylation quantitative trait loci – mQTL; +/- 1Mb). For CoRSIVs 1.0^2^, most mQTL effects were found to have a negative slope (i.e. the major allele was associated with higher methylation). Analysis of the 715 informative CoRSIVs 2.0 with significant mQTL likewise found that most of the mQTL effects have a negative slope (Fig. 3e, Supplementary Table S11). Evaluating the strength of mQTL effects, however, uncovered a substantial difference from CoRSIVs 1.0. In the original target-capture data^2^ the median mQTL R^2^ was 0.76, and only 16% of CoRSIVs yielded an mQTL R^2^<0.5. CoRSIVs 2.0, however, show a clear bimodal distribution, with about half showing strong mQTL (with a peak R^2^∼0.8) and the other half showing weak mQTL (with a peak R^2^∼0.1) (Fig. 3f). This finding is consistent with the elimination of contamination by CpG SNVs within CoRSIVs, genetic variation that was interpreted as epigenetic variation. Many CoRSIVs with weak mQTL nonetheless show strong SIV (Supplementary Fig. 8), suggesting they may be metastable epialleles^15–17^.

**Fig. 3.**
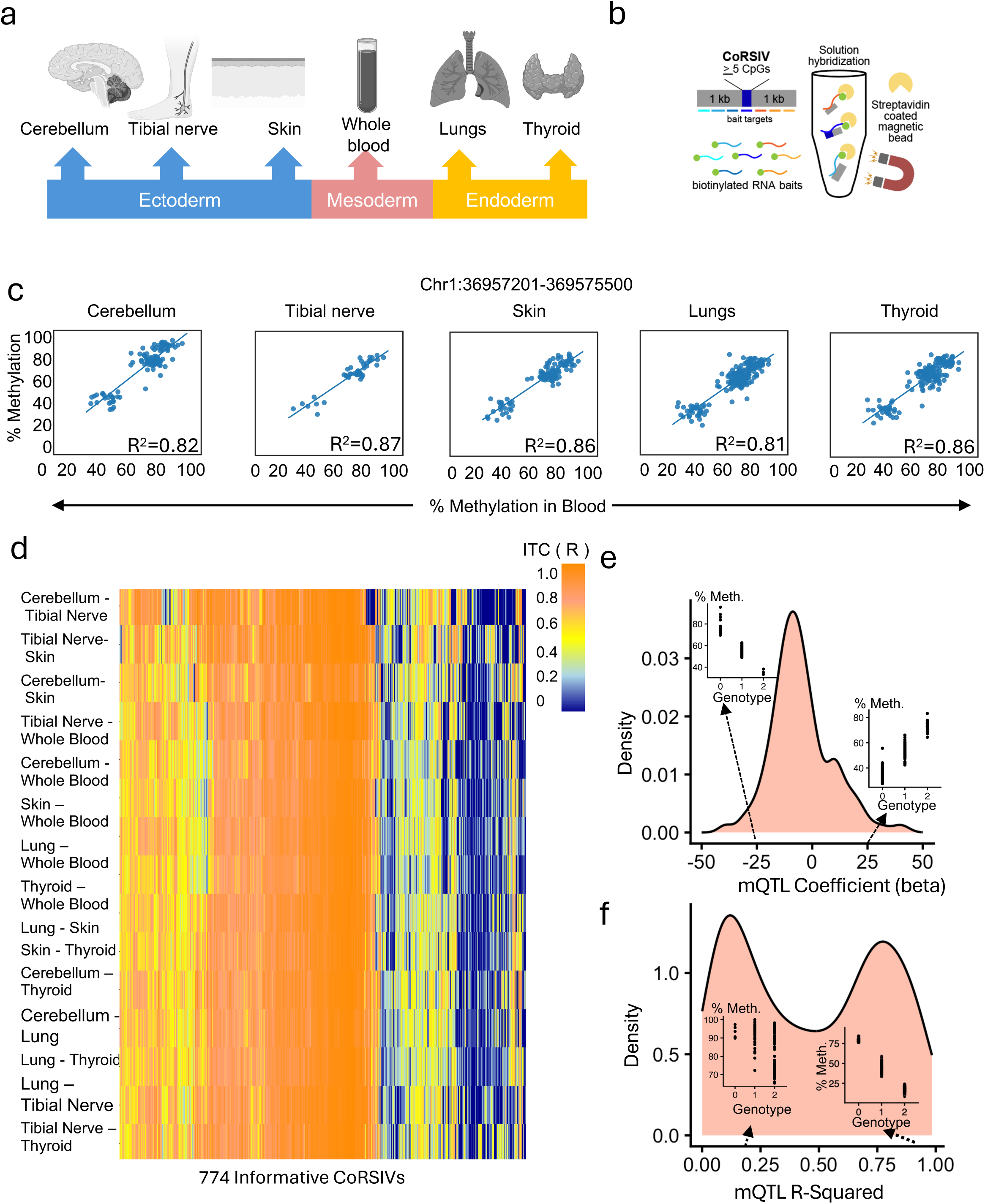
Validation of systemic interindividual variation and analysis of methylation quantitative trait loci (mQTL). **a** Genomic DNA samples were obtained from multiple tissues (representing the three embryonic germ layers) from each of up to 188 GTEx donors. **b** CoRSIV capture process using Agilent reagents. **c** At a representative CoRSIV, target-capture bisulfite sequencing data illustrate positive inter-tissue correlations for cerebellum, tibial nerve, skin, lung and thyroid vs. blood. **d** Heat map of inter-tissue correlations across 774 CoRSIVs included in the dataset; each column is a CoRSIV and each row one of 15 possible pairwise comparisons. Generally high positive inter-tissue correlations validate systemic interindividual variation. **e** Distribution of beta coefficients of significant Simes mQTL associations in blood at the 715 informative CoRSIVs showing significant mQTL. Insets show examples of negative and positive mQTL beta coefficients; SNV genotype indicates the number of copies of the minor allele. **f** Distribution of Simes mQTL R^2^ in blood across the 715 CoRSIVs. Insets show examples of CoRSIVs with low and high mQTL R-squared, respectively.

While the 10-20 individuals in each screen are not sufficient for a CoRSIV-wide mQTL analysis, we asked whether the greater number of CoRSIVs in Black versus White Americans might be explained by genetic variation in their vicinity. Since genetic variants that confer strong mQTL tend to be within 10kb of CoRSIVs^2^, we used 1000 Genomes data on Black and White Americans (ASW and CEU) to examine the density of common SNVs within 10kb of Black and White CoRSIVs, as well as both types of control regions. The results (Supplementary Fig. 9) show that CoRSIVs have the most common SNVs in their vicinity, followed by matched control regions (which have the same CpG density), then by random controls. As expected^18^, for each type of region, Black Americans have more common SNVs within 10kb than do White Americans. Curiously, however, whereas the DNA flanking matched and random control regions contains ∼40% more SNVs in Black than White Americans, in the vicinity of CoRSIVs this increment is only 26%. (Supplementary Fig. 9). Hence, the greater number of CoRSIVs in Black compared to White Americans is not straightforwardly attributable to their greater genetic diversity.

### Establishment of methylation at CoRSIVs 2.0 is sensitive to periconceptional environment

Multiple studies have documented effects of periconceptional environment on establishment of methylation at human genomic regions of SIV (sometimes dubbed ‘candidate metastable epialleles’), most notably in the context of season of conception in subsistence farmers in the Gambia, West Africa^3,4,9^. These effects were also documented in our initial report on human CoRSIVs^1^. Here, we used multiple data sources to evaluate a range of periconceptional influences on DNA methylation at CoRSIVs 2.0. We first explored effects of assisted reproductive technologies (ART); following *in vitro* fertilization, cleavage-stage human embryos are cultured for several days in synthetic media, a potent early embryonic exposure. We downloaded a public dataset with Illumina EPIC array methylation data on blood DNA of ART-conceived (N=193) and naturally conceived individuals (N=86)^19^. The same individuals were profiled at birth and in adulthood (age 18-28 y). Analyzing only data on the 3,017 EPIC probes within CoRSIVs 2.0 or matched control regions, we used DMRcate^20^ to identify differentially methylated regions (DMRs) between ART and naturally-conceived newborns. Of the 13 DMRs that met the DMRcate-recommended criteria of FDR < 0.05 and |Δβ| > 5%, all were within CoRSIVs (Fig. 4a) (Chi-sq Test: P=2.0x10^-298^). Analyzing the data from these same individuals in adulthood (Fig. 4b) detected 11 DMRs, again all within CoRSIVs (Chi-sq Test: P=4.1x10^-282^). Six DMRs were detected at both ages, and in the same direction (Fig. 4, a and b; Supplementary Table S12), indicating that, at CoRSIVs, DNA methylation alterations induced by early embryonic environment are stable over the life course. Accordingly, in a complementary approach, we asked whether probe-level CoRSIV methylation data from newborns can be used to train a machine-learning classifier to identify adults conceived by ART. Indeed, of various classifiers evaluated, a Support Vector Machine model trained on newborn CoRSIV methylation data achieved high sensitivity and specificity, with an area under the ROC curve (AUC) of 0.93 (Fig. 4c). These results indicate that influences of periconceptional environment on CoRSIV methylation persist to adulthood and provide the first evidence that adults conceived by ART carry a robust epigenetic signature of their *in vitro* origins. The effect of ART on establishment of methylation at CoRSIVs was corroborated in another dataset in which peripheral blood of newborns (94 ART and 43 naturally-conceived) was profiled by the Illumina HM450 array^21^. Applying the same approach as in the other data set detected 87 ART vs. natural-conception DMRs in CoRSIVs and only 8 in matched control regions (Fig. 4d) (Supplementary Table 13), confirming the high sensitivity of CoRSIVs to ART (Chi-sq Test; P=9.8x10^-110^).

**Fig. 4.**
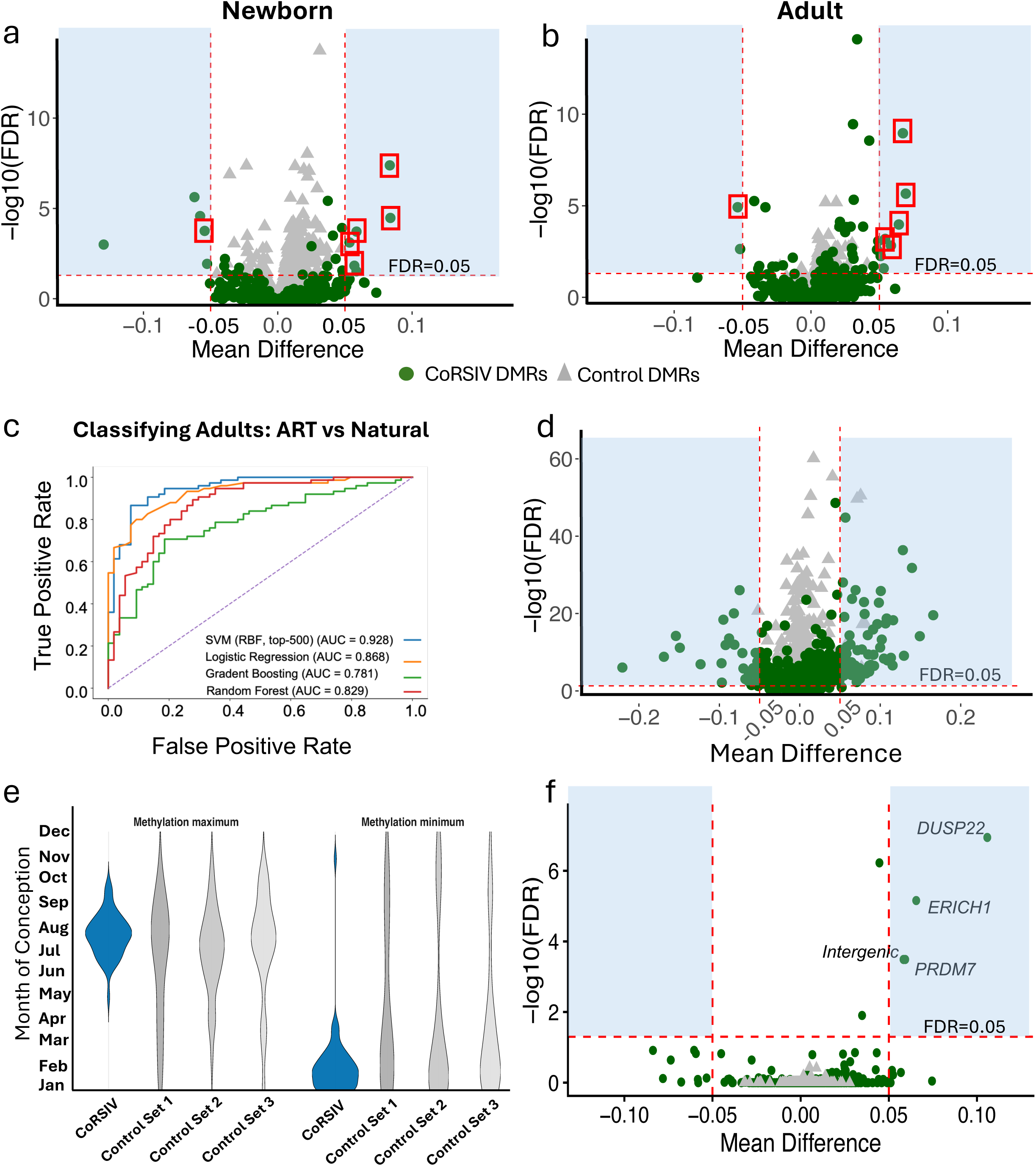
Diverse datasets highlight the sensitivity of CoRSIVs to periconceptional environment. **a** and **b** To detect methylation differences in blood of newborns (**a**) and adults (**b**) conceived by assisted reproductive technologies (ART, N=193) vs. naturally (N=86), Illumina EPIC array data^19^ within CoRSIVs (circles) and control regions (triangles) were used to identify differentially methylated regions (DMRs). All significant DMRs (shaded regions) were within CoRSIVs; of the 13 and 11 detected in newborns and adults, respectively, 6 detected at both ages are marked with squares. **c** Classification of adults as ART or naturally conceived. Of several machine-learning algorithms trained on the newborn CoRSIV data^19^ and tested on the adult CoRSIV data, support vector machines (SVM) performed best, achieving an AUC of 0.93. **d** An Illumina HM450 data set on 94 ART and 43 naturally-conceived newborns^21^ confirms the sensitivity of CoRSIVs to ART. Twelve times more DMRs were detected among CoRSIVs than in control regions (95 vs. 8). e) Consistent with previous reports^4^, establishment of methylation at CoRSIVs 2.0 is particularly sensitive to season of conception in 233 rural Gambian 2-year olds. f) Illumina HM450 data on blood of 75 adults who were exposed to the great Chinese famine during their first trimester of prenatal development, versus 105 unexposed controls^22^ corroborate CoRSIVs’ sensitivity to periconceptional nutrition. Among CoRSIVs, 4 significant DMRs (at *DUSP22*, *ERICH1*, *PRDM7*, and chr6:intergenic) show higher methylation in famine-exposed individuals. None was detected within control regions.

We next evaluated data on the effects of season of conception in a rural Gambian cohort of 233 two-year old children who were conceived throughout a calendar year (Fig. 4e). Here we observed increased methylation in individuals conceived in the Gambian rainy season (July to September) at CoRSIVS, replicating our previous finding in the same cohort at CoRSIVs 1.0^1^, with a greatly attenuated effect in matched control regions. As an additional level of scrutiny, we included two additional sets of matched controls, generated the same way as the first. Lastly, we analyzed a dataset designed to identify persistent epigenetic effects of prenatal exposure to the Great Chinese Famine of 1959-1961. The Illumina HM450 array was used to profile blood DNA methylation in 48 adults who had been exposed to famine during their first trimester of prenatal development, versus 105 unexposed controls^22^. Remarkably, though the subjects were ∼50 years old when studied, four significant DMRs were detected, all within CoRSIVs; these are associated with *DUSP22*, *ERICH1*, *PRDM7*, and one intergenic region (Fig. 4f) (Chi-sq Test: P=4.8x10^-144^). All 4 show higher methylation in individuals who had been exposed to famine prenatally (Supplementary Table 14). Only the *DUSP22* DMR was detected in the original publication^22^. Together, these results indicate that, across a wide range of exposures, settings, and ancestries, establishment of DNA methylation at CoRSIVs 2.0 is particularly sensitive to the environment of the pre-implantation embryo, and that these induced methylation changes persist over the life course. Analyzing the hundreds of thousands of CpG sites covered by the Illumina arrays entails a severe multiple testing penalty, compromising power. Re-analysis of existing data sets, targeting only the EPIC v1 or v2 array probes within CoRSIVs (Supplementary Tables S15 and S16, respectively) may thus reveal previously-undetected exposure- and disease-associated epigenetic variation^23^. The matching control EPIC probes used in this analysis are provided in Supplementary Table S17.

### Transcription factor binding data link CoRSIVs to development and disease

To evaluate the relevance of CoRSIVs to human health, we generated lists of genes associated with CoRSIVs 2.0 and performed gene set enrichment analysis using Enrichr^24^ to evaluate associations with the Jensen diseases and Disgenet databases. To conduct equally powered analyses on the Black and White CoRSIV sets we down-sampled from the list of genes associated with Black CoRSIVs to generate 10 random subsets of 1035 genes each (Supplementary Table S18) – matching the number of genes associated with White CoRSIVs – and averaged the results. This analysis (Fig. 5a) showed that both Black and White CoRSIVs are associated with cancer; liver, kidney, and breast cancer are top terms in both analyses. Black and White CoRSIVs are also associated with neurodevelopmental disorders including autism, schizophrenia, and attention-deficit/hyperactivity disorder. Notably, the Enrichr results converged on similar disease terms even though the genes associated with Black and White CoRSIVs are >75% different (Supplementary Fig. 10).

**Fig. 5.**
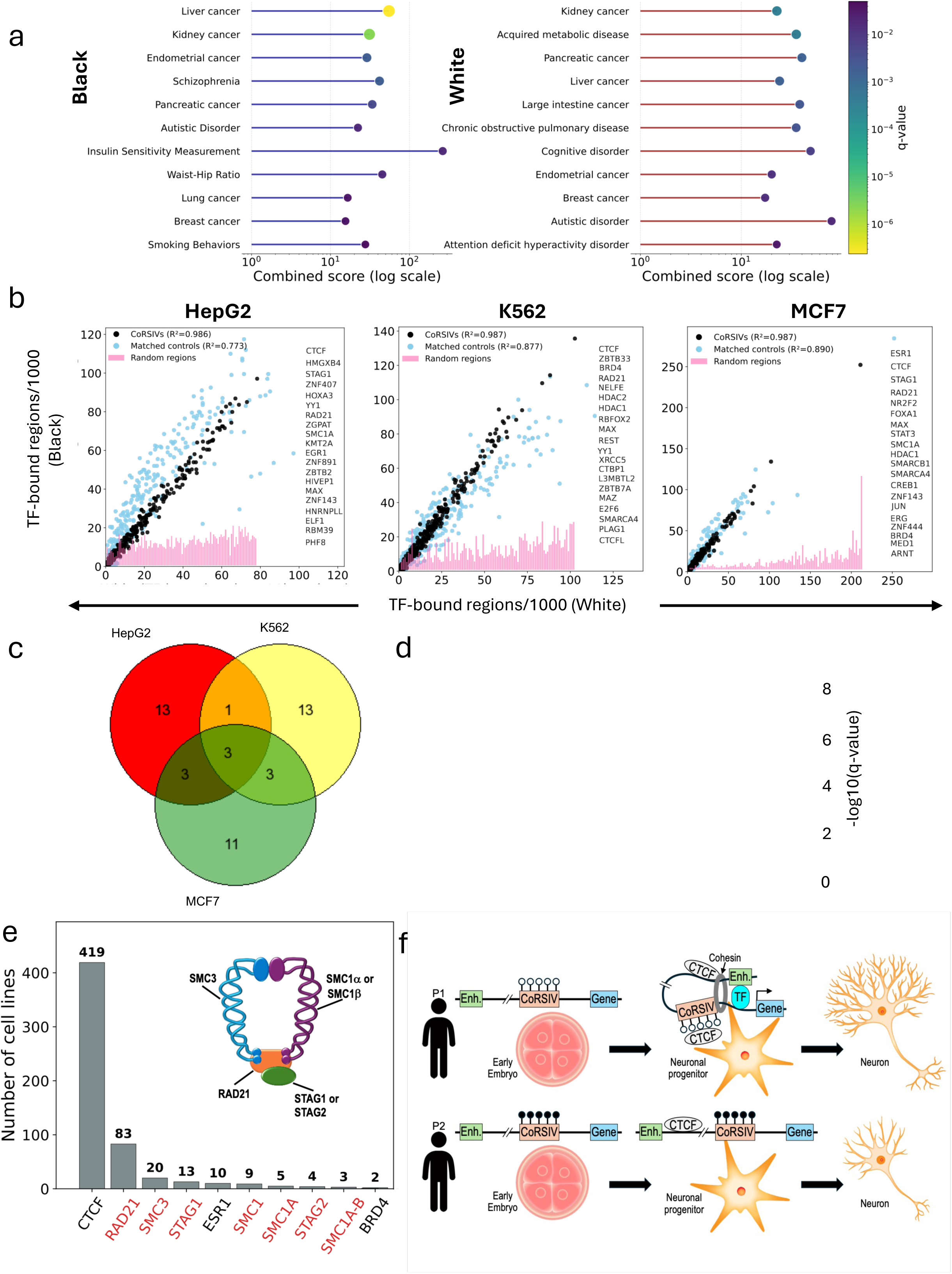
Association of CoRSIVs with diseases and transcription factor binding. **a** Gene set enrichment analysis shows that genes associated with Black (left) and White (right) CoRSIVs are associated with cancer and neurodevelopmental disorders. **b)** Analysis of transcription factor binding to CoRSIVs, based on ReMap. Scatter plots of normalized transcription factor overlap frequencies across White, Black, and matched control regions. For each cell line (HepG2, K562, MCF7), the top 100 transcription factors (TFs), ranked by summed normalized overlap counts across White CoRSIVs, Black CoRSIVs, and matched control regions, are shown. Each point represents one TF. Top 20 TFs are listed to the right of each plot. The x-axis shows the number of White CoRSIVs bound by that TF (per 1,000 regions), while the y-axis shows the corresponding normalized overlap frequency for either Black CoRSIVs (black points) or Black matched Controls (blue points). Normalized counts for these same TFs binding to random controls (genomic background) are shown in pink bar plots. **c)** Venn diagram of the top 20 TFs that bind CoRSIVs in each of the three cell lines. Most are specific to one cell line. **d)** Heatmap of disease-term enrichment for TFs associated with black and white CoRSIV sets in HepG2, K562, and MCF7 cells. **e)** Bar chart showing, for each TF, the number of Remap2022 biotypes in which it ranked among the top 5 most enriched factors (enrichment score ≥10). CTCF was the top-ranked factor in 419 biotypes; all four cohesin complex subunits (red labels) dominated the remainder of the top 10. Inset illustrates the four-subunit structure of cohesin. **f)** Model for how interindividual differences in CoRSIV methylation, established in the early embryo, regulate transcription factor binding and genome organization to modulate cellular differentiation, leading to different phenotypic outcomes.

To begin to understand how DNA methylation states established in the cleavage-stage embryo might influence later risk of disease, we analyzed data on transcription factor (TF) binding at CoRSIVs. We assessed the genomic intersections of Black and White CoRSIVs, and both types of control regions, with TF binding based on chromatin immunoprecipitation (ChIP-seq) peaks across three different human cancer cell lines (HepG2 (liver), K562 (leukemia), and MCF7 (breast)). These represent the most-studied human biotypes in Remap2022, a manually curated database encompassing 7,942 human ChIP-seq experiments^25^. For each target region, we identified all the TFs with evidence of binding to that region in each of the three cell lines. The raw counts of CoRSIVs with evidence of TF binding were normalized relative to the total number of regions in each set, yielding a standardized metric of TF-bound regions per 1,000 queried regions (Supplementary Table S19). We initially observed that within each of the three cell types the top TFs binding to Black and White CoRSIVs were nearly identical. Indeed, for each of the top 100 TFs binding to the largest number of White CoRSIVs, the proportion of Black CoRSIVs bound by the same TF is highly similar, yielding linear correlations with R^2^=0.99 in all three cell lines (Fig. 5b). This high correlation is surprising because only 15% of Black CoRSIVs overlap White CoRSIVs. We postulated that most of the TF binding might occur within these shared regions, but analysis of top TFs (Supplementary Fig. S11) shows that only 9-13% of TF-bound Black CoRSIVs overlap with TF-bound White CoRSIVs, similar to the overlap overall. To put these findings in context, we analyzed data on TF binding within both types of control regions. In all three cell lines, the top 100 TFs bound to white CoRSIVs bound to only 1/3 as many Black random control regions (Fig. 5b). TF binding to matched control regions, by comparison, showed overall levels comparable to CoRSIVs (Fig. 5b), indicating that CoRSIVs’ CpG density is a key driver of TF binding. Rather than dismissing the enhanced TF binding to CoRSIVs as merely a consequence of their CpG density, it is notable that their CpG density is very rare in the human genome. Genomewide, there is about 1 CpG per 100 bp; CpG islands, a major focus of the epigenetics field for the last 30 years, contain ∼8-12 CpGs per 100 bp (Supplementary Fig. S12). By comparison, 2/3 of CoRSIVs have an intermediate CpG density of 2-8 per 100 bp, an understudied category of the genome. Moreover, the coordination of binding proportions to Black vs. White matched control regions was substantially less than for CoRSIVs (R^2^=0.77,0.87, and 0.89, P= 4.1x10^-30^, 4.8x^-39^, and 9.1x10^-59^ in HepG2, K562, and MCF7, respectively). Of the top 20 TFs bound most commonly to White CoRSIVs, most are unique to one cell type (Fig. 5c), indicating that TFs bind to CoRSIVs in a highly cell type-specific manner. Nonetheless, except for the stronger association with cancer in K562 cells, gene set enrichment analysis based on the top 100 CoRSIV-binding TFs yielded similar enrichments for each of the three cell types (Fig. 5d). Enrichment of cancer-related terms in these cell lines is not surprising, but significant enrichments were also detected for intellectual disability and developmental syndromes related to neurological development, including Coffin-Siris and KBG. These similar enrichments were obtained although the genes targeted by the top 100 TFs are largely different for Black and White CoRSIVs (Supplementary Fig. S13). Hence, the results of gene set enrichment analysis based on cell type-specific TF binding to CoRSIVs (Fig. 5d) echo those based on CoRSIVs overall (Fig. 5a).

TFs often bind DNA sequence motifs in cooperative combinations, achieving both greater specificity and intensity of transcriptional activation^26^; such combinatorial control commonly involves combinations of 5-15 TFs binding to each regulatory region^27^. We postulated that the high coordination of ranked TF binding to CoRSIVs might reflect the stoichiometry of combinatorial binding. Focusing on the top 100 TFs bound to CoRSIVs and control regions in each cell type, we evaluated the multiplicity of TFs bound to each region (Supplementary Fig. 14a). Compared to random control regions, each CoRSIV is much less likely to be bound by only one or two different TFs. Rather, in all three cell types, compared to either random or matched control regions, each Black or White CoRSIV is *more* likely to be bound by 5-15 TFs (Welch T-test P<10^-16^) (Supplementary Fig. S14b). These findings indicate that CoRSIV methylation regulates combinatorial TF binding during development, providing a potential explanation for the highly coordinated hierarchy of TF binding to Black and White CoRSIVs (Fig. 5b).

As a complementary, approach to identify TFs most strongly associated with CoRSIVs, of the 7,942 human Chip-seq experiments in the ReMap database we identified 780 that show significant enrichment of TF binding to CoRSIVs relative to random control regions (Chi-sq test: P<10^-10^)(Supplementary Table S20). To integrate across the 7,942 experiments, we ranked TFs by enrichment significance within each of the 780 profiled cell lines to identify the top five most enriched TFs per cell line. This analysis (Fig. 5e) showed that CTCF is the TF most strongly enriched for binding to CoRSIVs. Strikingly, of the other nine TFs in the top ten, seven correspond to all four subunits of the cohesin complex (Fig. 5e). CTCF is a major regulator of development; one of its key roles is to function together with cohesin to stabilize DNA loops that form during differentiation and are a key determinant of genome organization^28–30^. By comparison, whereas CTCF was also most strongly enriched for binding to matched control regions and CpG islands, only one cohesin subunit – RAD21 – was among the top ten (Supplementary Fig. 15). DNA methylation influences binding of many of the top TFs that bind to CoRSIVs (Fig. 5b), including CTCF. Together, these findings suggest that interindividual variation in DNA methylation at CoRSIVs, established in the cleavage-stage embryo, functions to regulate genome organization and transcriptional regulation during subsequent organogenesis and differentiation (Fig. 5f).

Because it provides greater phenotypic diversity at the population-level, interindividual epigenetic variation that influences developmental outcomes may be under evolutionary selection^31^. We therefore evaluated Tajima’s D, a classic metric to test for evidence that specific DNA sequences are evolving under a non-random process. White CoRSIVs showed modestly elevated Tajima’s D values relative to matched controls and Black CoRSIVs (Supplementary Fig. 16a), suggesting enrichment for intermediate-frequency variants. Because European populations generally exhibit higher Tajima’s D values than African populations due to demographic history^32^, these results are more consistent with CoRSIVs being enriched for common regulatory variation than with strong locus-specific balancing selection. A pairwise evolutionary metric, the fixation index^33^ (F_ST_) ranges from 0 to 1, with 1 representing complete differentiation of two populations. F_ST_ values obtained from gnomAD v3.1 for AFR (African) and NFE (Non-Finnish European) populations more clearly distinguish CoRSIVs (Supplementary Fig. 16b). Specifically, F_ST_ values are slightly higher in Black and White CoRSIVs than in matched control regions. Given that CoRSIVs are defined by systemic inter-individual variation in DNA methylation, elevated F_ST_ is consistent with increased genetic or epigenetic divergence at these loci. Together, the weak Tajima’s D signals and F_ST_ values close to zero suggest CoRSIVs are unlikely to be under strong balancing selection. Rather, the largely dissimilar CoRSIVs in Black and White Americans may reflect recent evolutionarily diversification.

## Discussion

Compared to interindividual genetic variation, interindividual epigenetic variation has attracted relatively little attention. Systemic interindividual epigenetic variants are particularly important, however, because such states are established during early embryonic development and stable thereafter, propagating to every cell type and representing a distinct level of molecular individuality in addition to variation in DNA sequence. Here, in addition to refining our genome-wide CoRSIV screen to avoid the impact of genetic variation at CpG sites, we expanded it to include Black as well as White Americans. Several previous studies have profiled DNA methylation across the human genome to identify CpG sites that show differences in methylation across self-identified racial groups. Three used Illumina arrays to profile methylation in lymphoblastoid cell lines (LCLs) from different population groups^34–36^, each finding that most but not all population-associated methylation differences are explained by genetic variation. Studies in primary blood cells of different ancestry groups^12,37,38^ yielded similar results, indicating that approximately 30% of differential methylation between ancestry groups is not explained by genetic variation, supporting the hypothesis that stochastic and/or environmental factors not captured by ancestry are substantial determinants of DNA methylation^37^. Whereas these previous studies mapped out human genomic CpG sites showing population-differences in methylation levels, we are not aware of previous studies assessing population-differences in interindividual *variation* in DNA methylation, other than by the Illumina array platform^39^. Hence, our current results provide the first unbiased insights into systemic interindividual epigenetic variation in Black Americans.

Although this refined set of CoRSIVs is substantially different from our original report, many characteristics are consistent with our previous findings^1,2^, including high CoRSIV density in subtelomeric regions, the MHC locus, and the chromosome 20 pericentromeric region, association with chromHMM quiescent and heterochromatin states, long-range associations with the same specific transposable element subfamilies, and sensitivity to periconceptional environment. Our analysis of mQTL found both a consistency and a major difference from our previous results. As initially reported^2^, mQTL effects at CoRSIVs 2.0 tend to have a negative coefficient, meaning the major allele is associated with higher CoRSIV methylation. But unlike the predominantly strong mQTL we reported earlier^2^, for CoRSIVs 2.0 the mQTL R^2^ distribution is strongly bimodal, with nearly half showing an mQTL R^2^ <0.5. This reduced magnitude of mQTL likely reflects the impact of masking genetically variant CpG sites, meaning that many of the associations originally interpreted as mQTL reflected haplotype effects. Future studies profiling a greater proportion of CoRSIVs 2.0 will be necessary to better understand the basis for the bimodal mQTL distribution. Nonetheless, our results suggest there may be many more metastable epialleles in humans than we previously thought, introducing a substantial novel source of molecular individuality not ‘hard-wired’ in the genome. This conclusion is consistent with previous studies^35,37,39^ that documented a substantial fraction of DNA methylation differences between human populations that is independent of genetic variation.

Our most surprising finding was the discovery of twice as many CoRSIVs in Black than White Americans. Whereas overall genetic diversity is not twice as high among Black than White Americans^40^, we postulated that perhaps CoRSIVs occur in hot spots of genetic variation among Black Americans. Our analysis of SNV density in the vicinity of Black and White CoRSIVs, however, did not support this. In the 10kb regions flanking matched and random control regions there are ∼40% more SNVs in Black than White Americans (Supplementary Fig. 9). In the 10kb regions flanking CoRSIVs, however, the excess of cis genetic variation in Black vs. White Americans was only 26%, contrary to our expectation. Hence, the much greater number of CoRSIVs in Black than White Americans is not easily explained by interindividual genetic variation. This finding is consistent with our mQTL analysis indicating that interindividual epigenetic variation at many human CoRSIVs is not determined by cis genetic variation (Figure 3f). The greater number of CoRSIVs in Black vs. White Americans despite the highly similar characteristics of CoRSIV-flanking genomic regions in these two racial groups is puzzling, both raising the question of what drives SIV and suggesting that substantially different genomic regions may exhibit SIV in different ancestry groups around the world. Indeed, whereas previous studies found that only 10-30% of CpG sites show methylation differences across ancestry groups^12,34–38^, most Black and White CoRSIVs are distinct. This indicates that, compared to average methylation, interindividual *variation* in DNA methylation shows greater differences amongst populations, consistent with the idea that genetic variants that increase interindividual epigenetic variation may be advantageous to evolutionary fitness^41^.

Consistent with our earlier reports on SIV regions^1,3,4^, the influence of periconceptional environment has a major impact on establishment of methylation at CoRSIVs 2.0. Many studies have documented altered DNA methylation in offspring conceived by ART. But a recent systematic review of 17 genome-scale studies^42^ found very few methylation alterations that replicated across studies, leading the authors to conclude that persistent epigenetic changes due to ART are minimal. The authors of the first ART dataset we analyzed^19^ likewise concluded that ART-associated epigenetic variation at birth largely resolves by adulthood. By focusing on the 3,017 EPIC probes within CoRSIVs, however, we found that most of the DMRs identified at birth persist to adulthood. Moreover, the stability of these induced methylation differences allowed us to show, for the first time, that CpG-level DNA methylation patterns in blood can be used to identify adults who were conceived by ART. In addition to documenting effects of ART, we confirmed previous reports^1,3,4^ that CoRSIV methylation is sensitive to season of conception in rural Gambian villagers. Lastly, our CoRSIV-focused re-analysis of EPIC array data on blood of Chinese adults^22^ (age ∼48) not only confirmed the reported persistent influence of early gestational famine exposure at *DUSP22* but identified three additional DMRs associated with early famine exposure. Together, these results indicate that establishment of CoRSIV methylation is sensitive to the environment of the early embryo, and that such environmentally-induced epigenetic alterations persist to adulthood. Moreover, although our CoRSIV 2.0 screen included only White and Black Americans, the data from the Chinese Famine cohort (Fig. 3f) indicate that many of these regions are also CoRSIVs in other ethnic and ancestry groups.

Analyses of gene set enrichment showed that both Black and White CoRSIVs 2.0 are associated with cancer and neurodevelopmental disorders. By comparison, CoRSIVs 1.0 were associated with metabolic and hematologic disorders, but not cancer^2^. To address how CoRSIV methylation states established in the early embryo may affect later disease outcomes, we hypothesized that interindividual epigenetic variation at CoRSIVs may modulate the binding of methylation-sensitive DNA binding proteins. Our analysis of ReMap ChIP-seq data (Figure 5b-e) supports this. Compared to genome background, CoRSIVs are strongly enriched for binding by important regulatory transcription factors and chromatin modifying complexes. Most striking, across the three most widely-studied ReMap biotypes, we discovered that the relative ranking of TF binding to Black and White CoRSIVs is nearly identical (R^2^=0.99). To some extent, the highly similar hierarchy of DNA binding proteins in Black and White CoRSIVs is likely driven by the cooperative nature many of these proteins. For example, HDAC1, HDAC2, and CTBP1, top DNA binding proteins in K562, form a core repressive complex^43^ regulating chromatin compaction, and SMARCA4 and SMARCB1, top DNA binding proteins in MCF7, are key components of the SWI/SNF complex^44^ that functions to reposition nucleosomes for chromatin remodeling. Despite this coordinated hierarchy of TF binding to CoRSIVs in Black and White Americans, however, the genes targeted by the TFs are largely different. CoRSIVs are genomic regions at which interindividual epigenetic variation is established prior to cellular differentiation. One interpretation, therefore, is that interindividual variation in developmental outcomes is evolutionarily advantageous, but individuals of African and European ancestry evolved different approaches to achieve it.

We have proposed^7^ that their systemic interindividual variation make CoRSIVs particularly advantageous for population study. Together, our latest data indicate that, much more, CoRSIVs may be key developmental regulators. Accordingly, one’s CoRSIV methylation profile, detectable in blood, may represent an epigenetic code providing information about the molecular instructions that guided his or her development. Together with the extensive evidence that establishment of CoRSIV methylation is influenced by the environment of the early embryo, our results suggest that CoRSIV methylation provides a mechanistic link between periconceptional environment, cellular differentiation, and phenotypic outcomes.

## Methods

### Conducting the CoRSIV screen

For the 10 White GTEx donors, we retrieved all SNVs with minor allele frequency (MAF) ≥ 0.05 in the 1000 Genomes CEU population (Utah residents (CEPH) with Northern and Western European ancestry). Of these, we retained only SNVs 1) that affect either a C or G at a CpG site, and 2) for which at least one of the 10 individuals in the screen carries the alternate allele. All of these affected CpG sites were masked in the coverage files of all 10 individuals. For the 10 Black GTEx donors, we retrieved all SNVs with MAF ≥ 0.05 in the 1000 Genomes ASW population (African Ancestry in Southwest US). Of these, we retained only SNVs 1) that affect either a C or G at a CpG site, and 2) for which at least one of the 10 individuals in the screen carries the alternate allele. All of these affected CpG sites were masked in the coverage files of all 10 individuals. Genome-wide identification of CoRSIVs followed the previously described two-stage approach^1^. Briefly, neighboring 100-bp bins containing informative CpGs were first aggregated into blocks of correlated methylation using pairwise Pearson correlation thresholds. For all such blocks, systemic interindividual variation (SIV) was then evaluated by assessing inter-tissue correlations (ITC) across matched tissues: brain vs. heart, thyroid vs. heart, and thyroid vs. brain (minimum ITC cutoff R≥0.71, R^2^≥0.5). This minimum ITC of 0.71 is a reliable standard for identifying SIV^15^. Regions meeting predefined thresholds^1^ for CpG content (≥5 CpGs) and interindividual average methylation range (≥20%) were retained for downstream analyses.

### Selecting matched control regions

CoRSIVs consist of contiguous 100 bp CpG bins, with adjacent bins separated by no more than 100 bp. To enable matched control selection, we first preprocessed all genomic 100 bp CpG bins. For each chromosome c and bin b, we computed the maximum length of a contiguous cluster of successive bins beginning at b, subject to the same ≤100 bp spacing constraint. For each CoRSIV located on chromosome c, comprising n CpG bins, genomic span s, and total CpG count g, we selected a matched control region by restricting to chromosome c and randomly sampling a bin b that initiates a cluster of n successive bins under the same spacing criteria. Control regions were required to match CoRSIVs within a target relative error e = 10% for both genomic span and CpG content. Specifically, we required that (i) the total span approximates s within e%, and (ii) the total CpG count approximates g within e%.

### Selecting random control regions

To establish a simple null model approximating the genomic background, we generated random control regions by random sampling under minimal constraints. Specifically, we fixed the length of control regions to the median genomic span of the CoRSIVs, thereby preserving the first-order length distribution while randomizing all other genomic features. These regions were then sampled uniformly at random from the genome, without conditioning on chromosome, CpG density, or local sequence context. This corresponds to sampling from a uniform distribution over all possible genomic intervals of the specified length.

### Genomic enrichment analyses

For enrichment analyses involving genomic annotations, including subtelomeric regions, candidate cis regulatory elements, promoters, pseudogenes, ChromHMM states, and other feature classes, odds ratios were calculated from contingency tables. Statistical significance was assessed using Pearson’s χ² test or, when expected cell counts were small, Fisher’s exact test.

### Correlation and distribution analyses

Associations between continuous variables were evaluated using Pearson correlation coefficients. Comparisons of distributions were performed using non-parametric tests, including the Wilcoxon rank-sum test or Kolmogorov–Smirnov test, as appropriate. These approaches were used for analyses such as inter-tissue methylation concordance, transcription factor overlap concordance, and comparisons of SNV density or mQTL effect-size distributions.

### mQTL analysis

Cis-mQTL analyses were conducted as previously described^2^, to test for associations between genetic variation and DNA methylation at CoRSIVs using a locus-level inference framework based on the Simes procedure^45^. Genotype data for GTEx donors were restricted to common SNVs (minor allele frequency ≥5%). Analyses were conducted using data on blood DNA, including only CoRSIVs with methylation and genotype data available for at least 20 donors. DNA methylation at each CoRSIV was summarized as the mean across constituent CpGs and transformed to M-values prior to analysis. For each CoRSIV, associations with all variants located within ±1 Mb were assessed using rank-based correlation. Resulting p-values were combined using the Simes method to generate a single locus-level statistic for each CoRSIV. Multiple testing across loci was controlled using false discovery rate (FDR) correction within each tissue, with significance defined as FDR < 0.05. For CoRSIVs exhibiting significant mQTL signals, effect sizes were estimated using linear models, and the proportion of variance in methylation explained by genotype (R²) was calculated.

### Differential methylation analyses

Differentially methylated regions associated with assisted reproductive technology, seasonal variation, or famine exposure were identified using DMRcate. Significance was defined using the recommended thresholds of false discovery rate (FDR) < 0.05 and absolute methylation difference (|Δβ|) > 5%. Multiple-testing correction was performed using the Benjamini–Hochberg procedure.

### Machine Learning Classification of ART vs. Natural Conception

We developed a machine learning framework that classifies individuals as ART-conceived or naturally conceived based on DNA methylation at Correlated Regions of Systemic Interindividual Variation (CoRSIVs). Classifiers were trained exclusively on neonate blood methylation data^19^ and tested on the same individuals in adulthood, providing a cross-life-stage test of epigenetic persistence. Prior to classification, methylation beta values were corrected for inter-individual variation in blood cell composition using OLS residualization against EpiDISH-estimated leukocyte fractions. Five supervised binary classifiers — Logistic Regression, Random Forest, Gradient Boosting, Support Vector Machine (RBF kernel), and XGBoost — were trained using 3,017 CoRSIV CpG features, with model performance assessed by ten-fold stratified cross-validation in neonates and AUROC on the held-out adult test set. All classifiers were trained exclusively on neonate samples (79 ART, 58 non-ART). The adult cohort (129; 54 ART, 75 non-ART) was held out entirely as an independent test set and was not used in any aspect of model fitting or hyperparameter selection.

### Gene set enrichment analyses

Genes are annotated to CoRSIVs based on genomic distance. If a CoRSIV is located +/-3kb of a transcription start site (TSS) or transcription end site (TES), or overlaps a gene body, it is annotated to that gene. For each group, the gene set is the aggregate of all annotated genes. Gene set enrichment analyses were performed using Enrichr^24^. Statistical significance was assessed using adjusted p-values reported by the platform, with emphasis on false-discovery-adjusted q-values. To account for differences in the number of CoRSIV-associated genes between Black and White donor sets, enrichment analyses for Black CoRSIV-associated genes were performed on size-matched random subsets, matching the number of genes associated with White CoRSIVs, prior to comparison.

### Comparisons of CpG density

CpG density was quantified for all CoRSIV 100-bp windows, a size-matched random genomic background, cohort-specific matched controls (Black and White populations separately), and annotated hg38 CpG islands (UCSC Genome Browser). For each region set, bedtools nuc (v2.x) was run against the hg38 reference genome using the -pattern CG flag to count CG dinucleotides within each interval. CpG density was expressed as the number of CpG sites per 100 bp (raw CG count divided by interval length, multiplied by 100). To compare the distribution of CpG density across groups, kernel density estimates (KDE) were plotted in Python using seaborn.

### Transcription factor binding analyses

Data on localization of transcription factor binding were obtained from ReMap2022^25^. To compare transcription factor occupancy across region sets of different sizes, the number of unique query regions overlapping at least one binding peak for a given transcription factor was normalized as overlaps per 1,000 queried regions. Concordance in normalized transcription factor occupancy between Black and White CoRSIVs was assessed using Pearson correlation coefficients.

### Unsupervised clustering and principal component analysis

Hierarchical clustering and principal component analysis were used to evaluate relationships among samples based on methylation data from CoRSIVs or matched control regions. Hierarchical clustering was performed using Euclidean distance and complete linkage unless otherwise stated. PCA was used as a descriptive dimensionality-reduction approach to visualize potential sample grouping by tissue, individual, or race.

### General statistical considerations

All statistical analyses were performed in R unless otherwise stated. All tests were two-sided. Multiple-testing correction was applied where appropriate, and adjusted p-values or q-values are reported in the corresponding figures, tables, or supplementary materials.

## Data availability

The whole-genome bisulfite sequencing (WGBS) datasets generated and analyzed in this study are available through the database of Genotypes and Phenotypes (dbGaP) under controlled access and can be accessed through the NHGRI Analysis, Visualization, and Informatics Lab-space (AnVIL) platform. Access requires dbGaP authorization and compliance with applicable data use agreements. The datasets are available under dbGaP accession phs001746: phs001746.v1.p1: WGBS data from 10 White GTEx donors used for the original CoRSIV discovery study. phs001746.v2.p1: CoRSIV v1 target-capture bisulfite sequencing data from 188 GTEx donors. phs001746.v3.p1(https://www.ncbi.nlm.nih.gov/projects/gapprev/gap/cgi-bin/study.cgi?study_id=phs001746.v3.p1): WGBS data from 10 Black GTEx donors generated in the present study. Previously published datasets used in this study were obtained from the Gene Expression Omnibus (GEO) under accession numbers GSE131433 and GSE79257.

## Code availability

All custom code used for data processing, CoRSIV discovery, statistical analyses, and figure generation is publicly available at the Waterland Laboratory GitHub repository (CoRSIV2.0): https://github.com/waterlandlab/CoRSIV2.0. The repository contains scripts, workflows, and documentation sufficient to reproduce the analyses reported in this study.

## Acknowledgements

We gratefully acknowledge the assistance of Dr. Kristin Ardlie at the Broad Institute, who facilitated our access to the GTEx samples, and Mr. Adam Gillum at Baylor College of Medicine, who helped to develop the figures. Funding for this project was provided by NIH/NIDDK (1R01DK129265) and the USDA/ARS (CRIS 3092-51000-065-003S). The Functional Genomics core at Baylor College of Medicine, where the WGBS data were generated, is partially supported by NIH shared Instrument grant S10OD023469. The GTEx Project was supported by the Common Fund of the Office of the Director of the National Institutes of Health, and by NCI, NHGRI, NHLBI, NIDA, NIMH, and NINDS.

## Author contributions

RAW conceived the study and obtained funding. CJG and WJC performed data analysis under the guidance of RAW. YL and RC oversaw production of the WGBS data. MSB managed laboratory aspects of sample handling. PI conducted data analysis under the supervision of MJS. MJS and AMP provided data for the Gambian season of conception analyses. GJ provided guidance on the analyses related to evolutionary selection. GH provided guidance on heritability and assessing influences of genetic variation. CC provided scripts and guidance to support the analyses of ChromHMM and transposable element associations with CoRSIVs. CJG and RAW wrote the manuscript with input from all coauthors. All authors read and approved the final manuscript.

---

**Figure S1.**
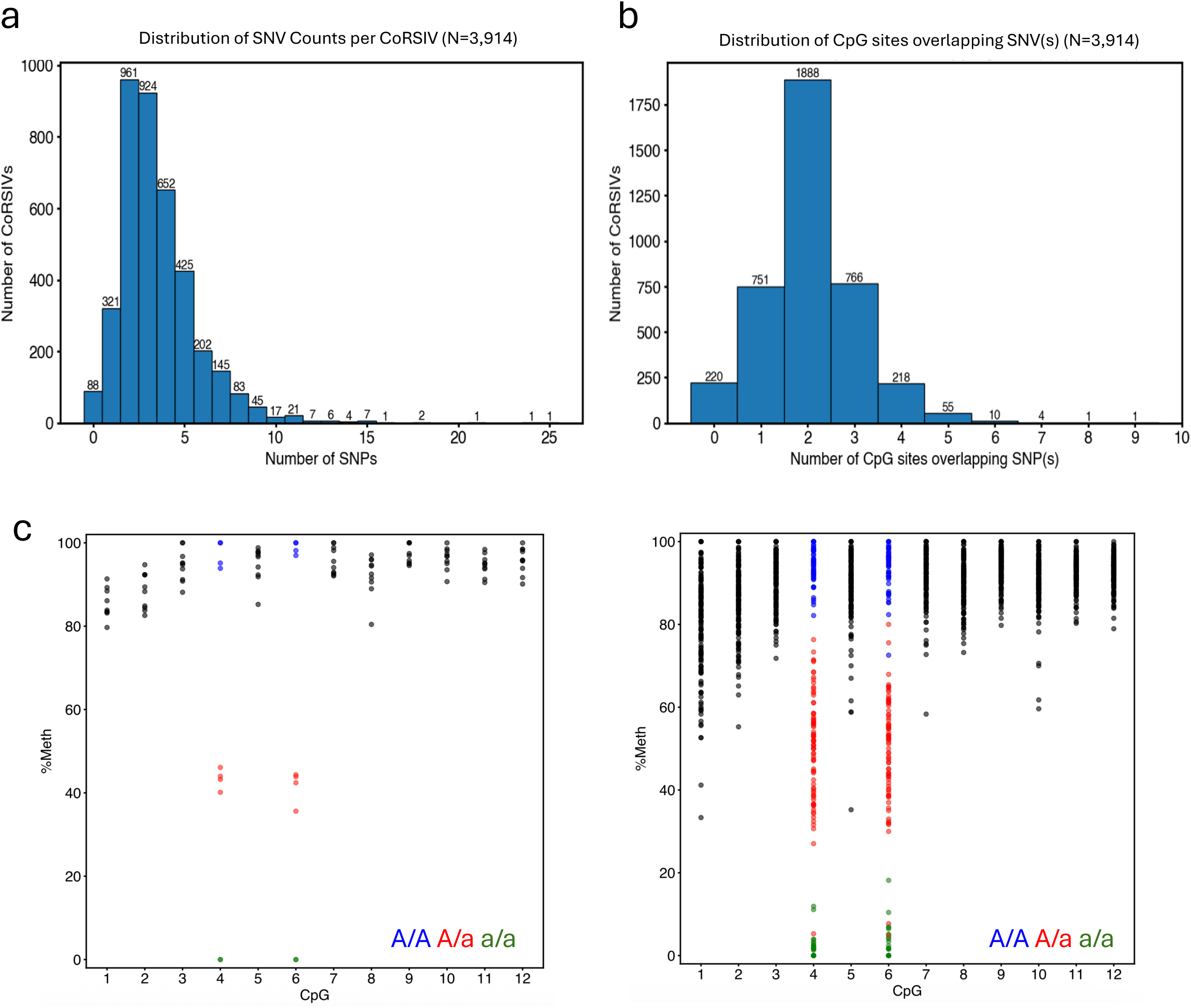
Effect of CpG-SNVs on CoRSIVs 1.0. **a)** Distribution of the number of SNVs within each of 3,914 CoRSIVs 1.0. **b)** Distribution of the number of CpG sites directly overlapping a SNV, within each of 3,914 CoRSIVs 1.0. **c)** Two examples in which much of the interindividual variation that was interpreted as methylation was due to genetic variation at SNVs. Each point is one individual. Genotypes at CpG SNVs are indicated by color: A/A (Blue) – Homozygous major allele, A/a (red) – Heterozygous, a/a (green) – Homozygous minor allele.

**Figure S2.**
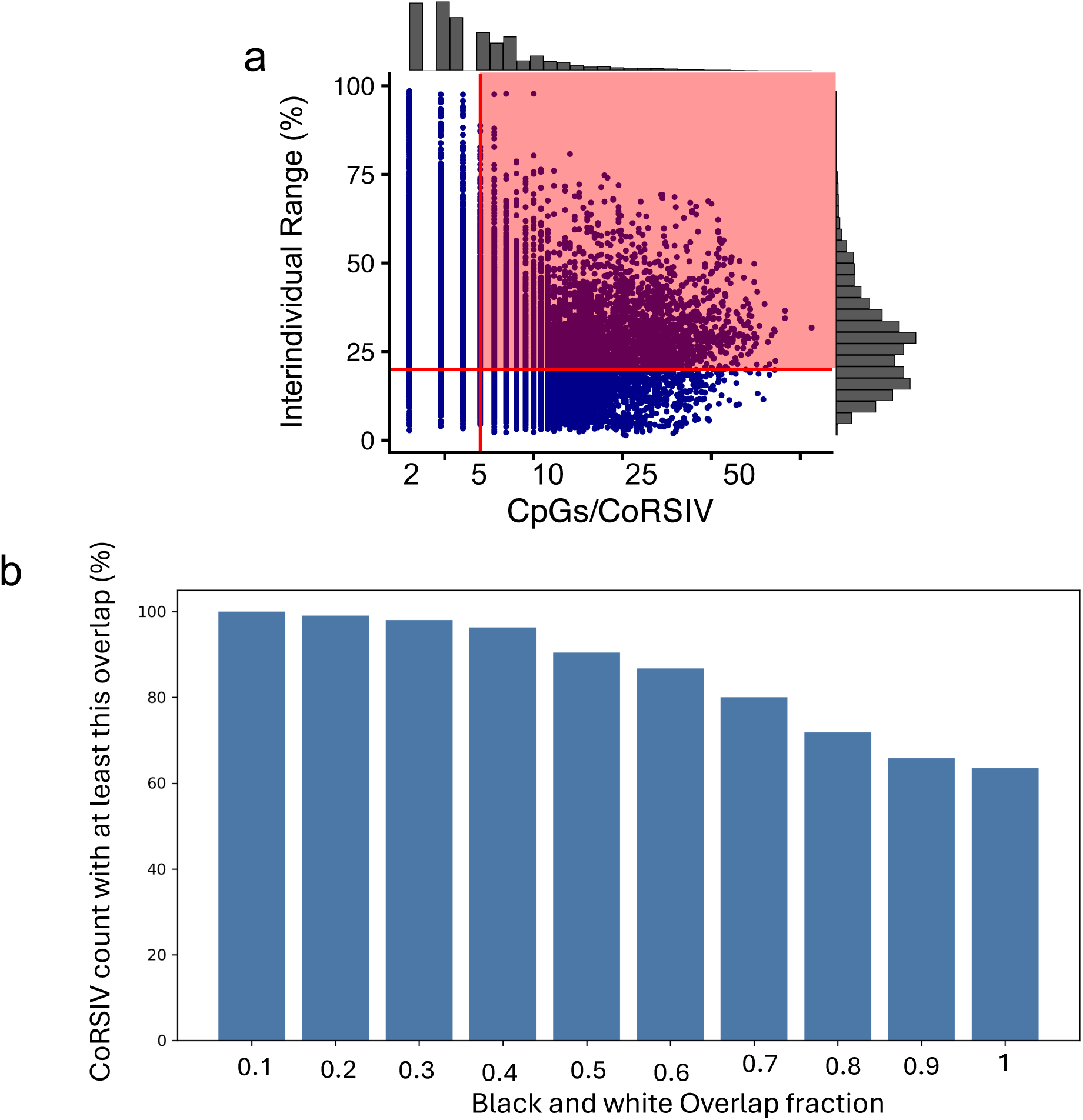
CoRSIV filtering and Black and White CoRSIV overlap. **a)** For the 39,368 CoRSIVs initially identified, scatter plot shows interindividual methylation range vs. number of CpGs per CoRSIV. Subsequent analyses focus on the 6,467 CoRSIVs each containing ≥ 5 CpGs and exhibiting an interindividual range ≥ 20% methylation (pink shaded area). **b)** Distribution of proportional overlap of all Black and White CoRSIVs that overlap at all; 63% of the CoRSIVs that overlap at all overlap completely (overlap fraction 1.0).

**Figure S3.**
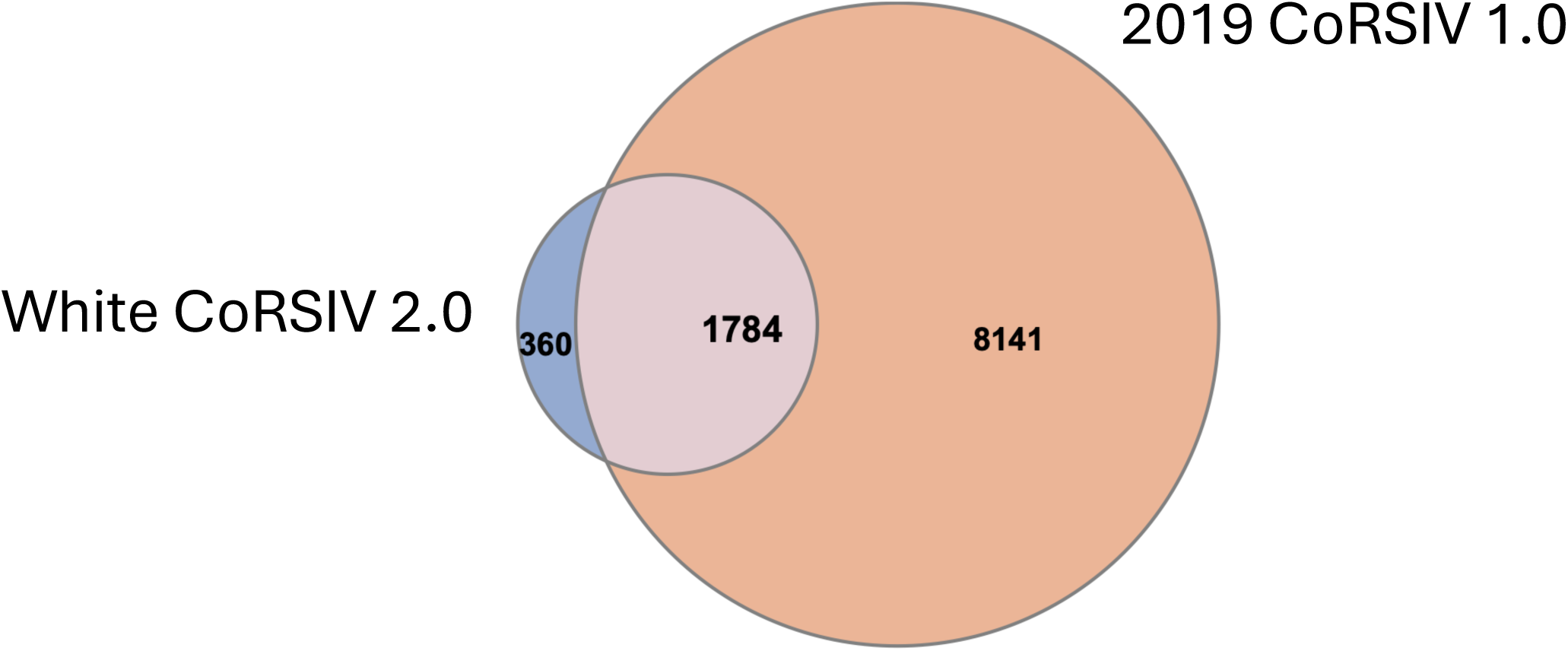
Genomic overlap (at least 1bp) between White CoRSIV 2.0 and CoRSIV 1.0.

**Figure S4.**
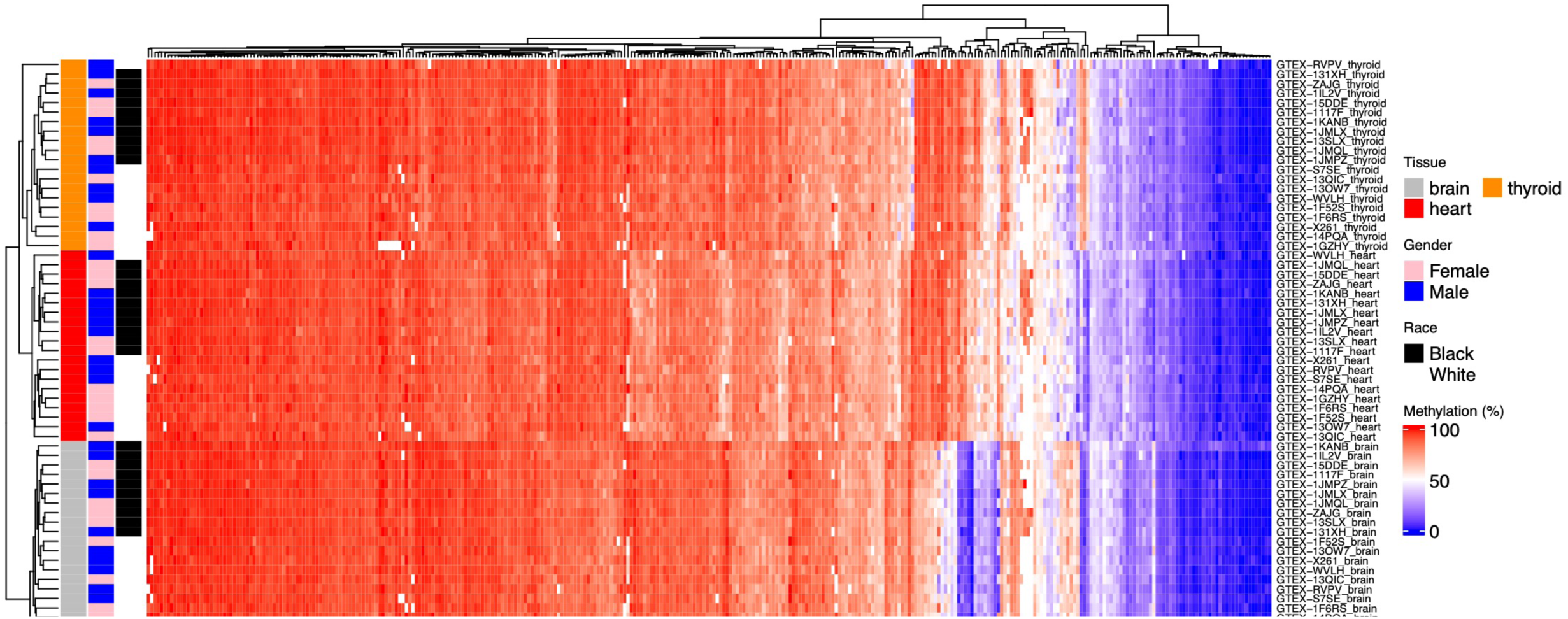
Unsupervised hierarchical clustering of DNA methylation in matched control regions. Unlike CoRSIVs, methylation data on matched control regions clusters samples by tissue, and nearly perfectly by race.

**Figure S5.**
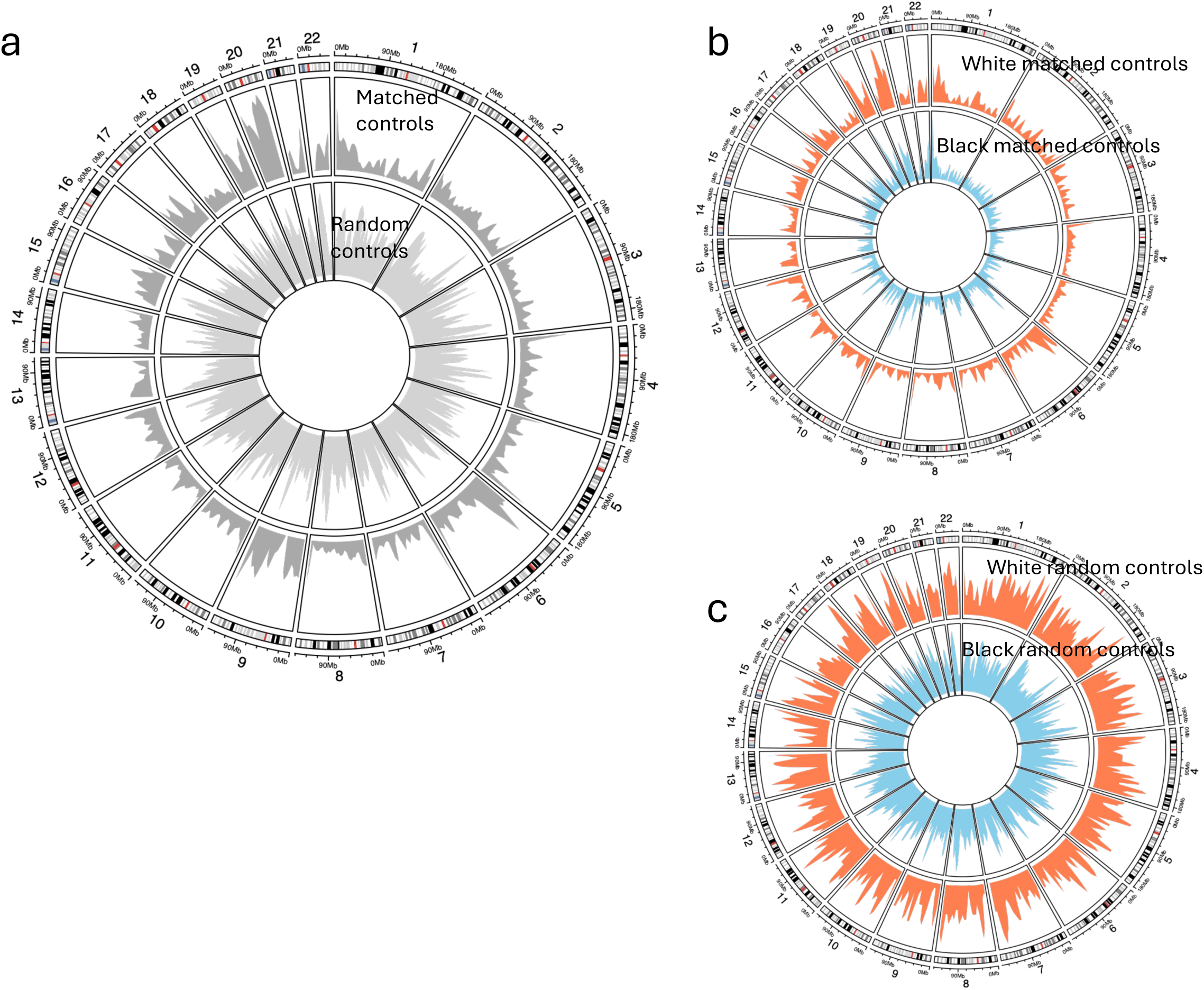
Genomic distributions of control regions. **a)**. The genomic distribution of matched control regions (outer ring) suggests that the high CoRSIV density in the MHC locus (chr. 6) (Fig. 2a) may be largely explained by CpG density. The high density of CoRSIVs in the pericentromeric region of chr. 20 (Fig. 2a) does not appear to be explained by CpG density. **b)** The distributions of control regions matched to Black (inner) and White (outer) CoRSIVs. **c)** The distributions of random control regions generated for Black (inner) and White (outer) CoRSIVs.

**Figure S6.**
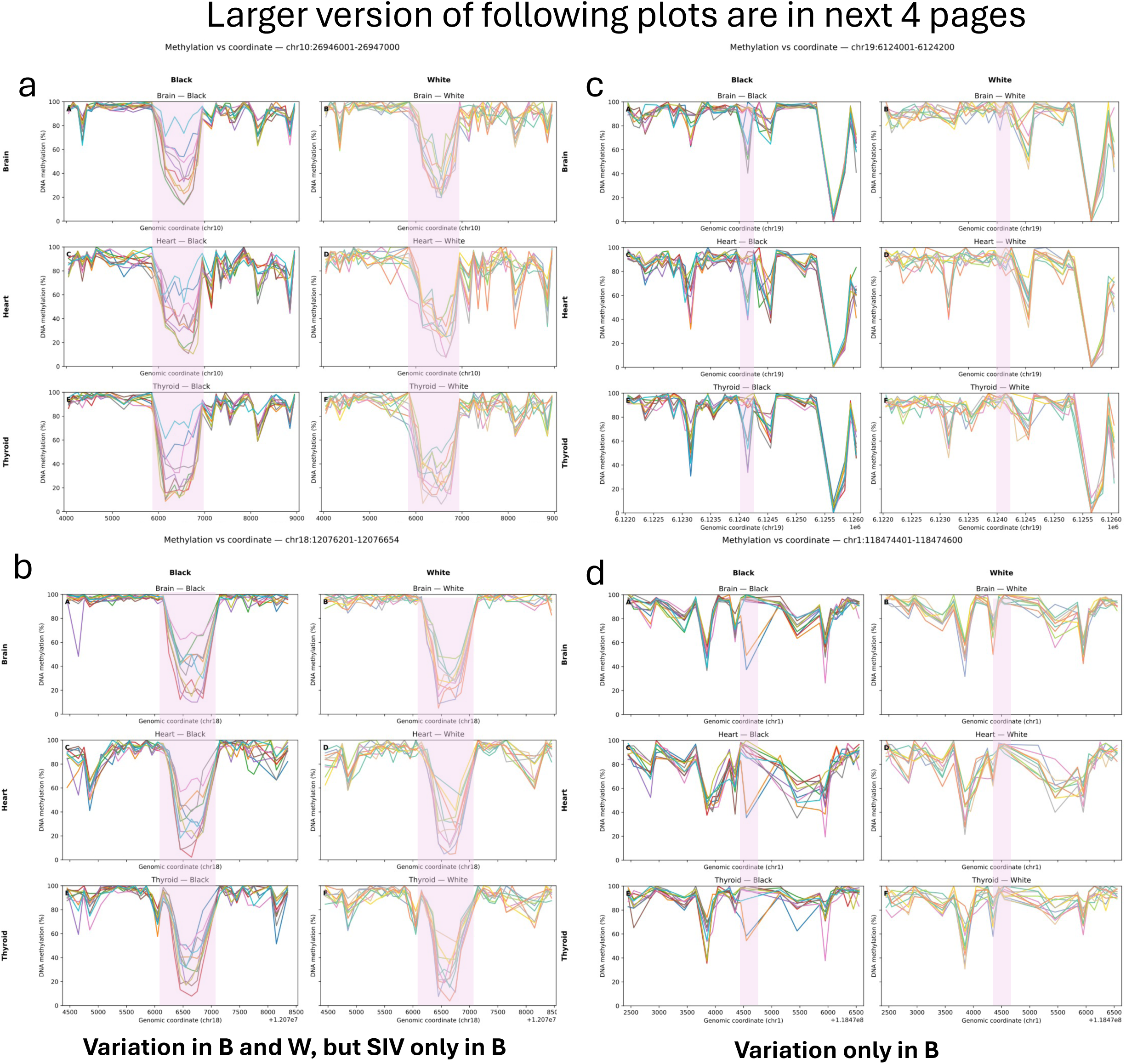
Examples of individual-level WGBS data for regions that are CoRSIVs among Black but not White donors. a,b) SIV among Black donors only; White donors show variation but no SIV (i.e. not systemic). **c,d)** SIV among Black donors only, and little variation among White donors.

**Figure S7.**
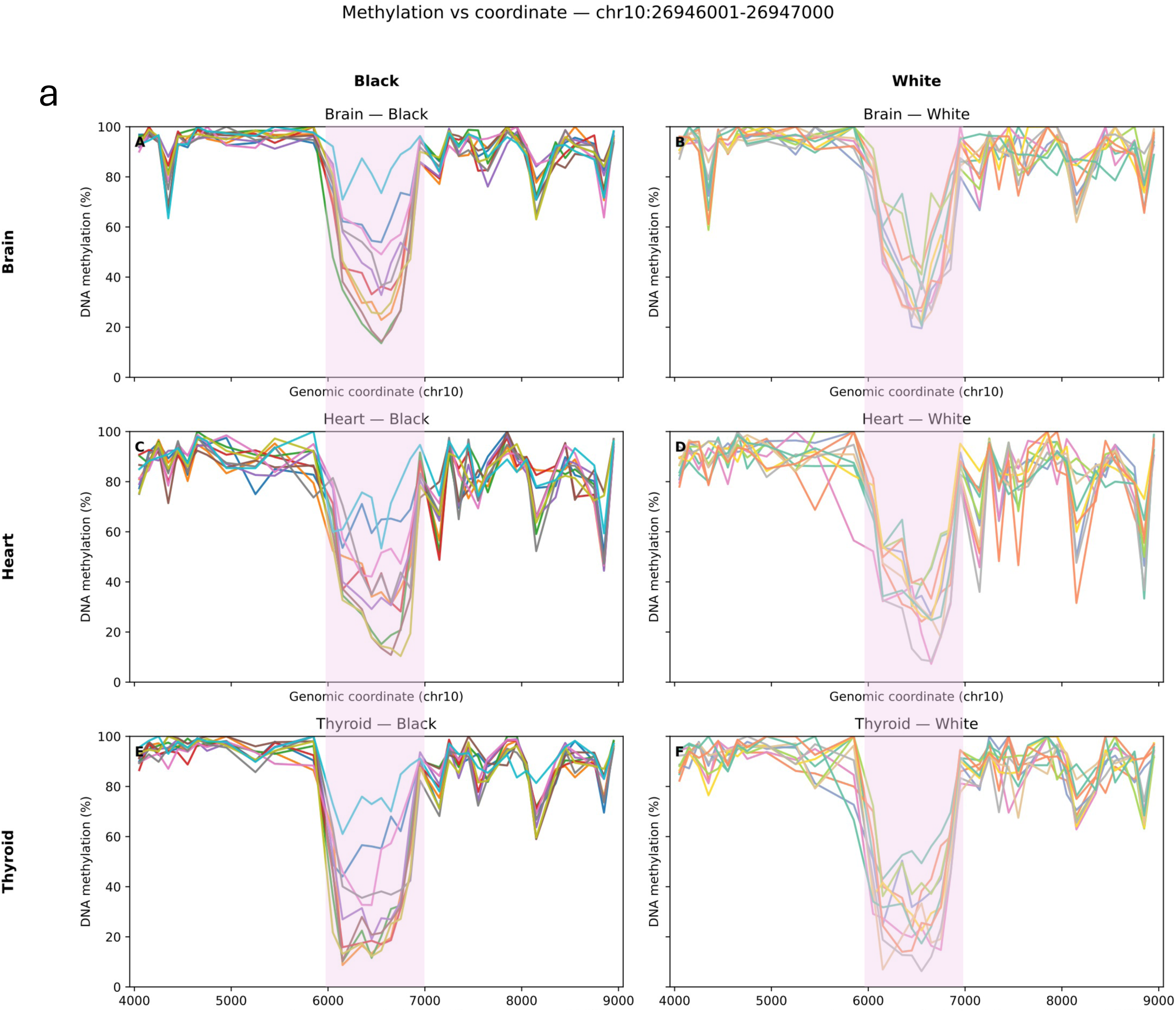
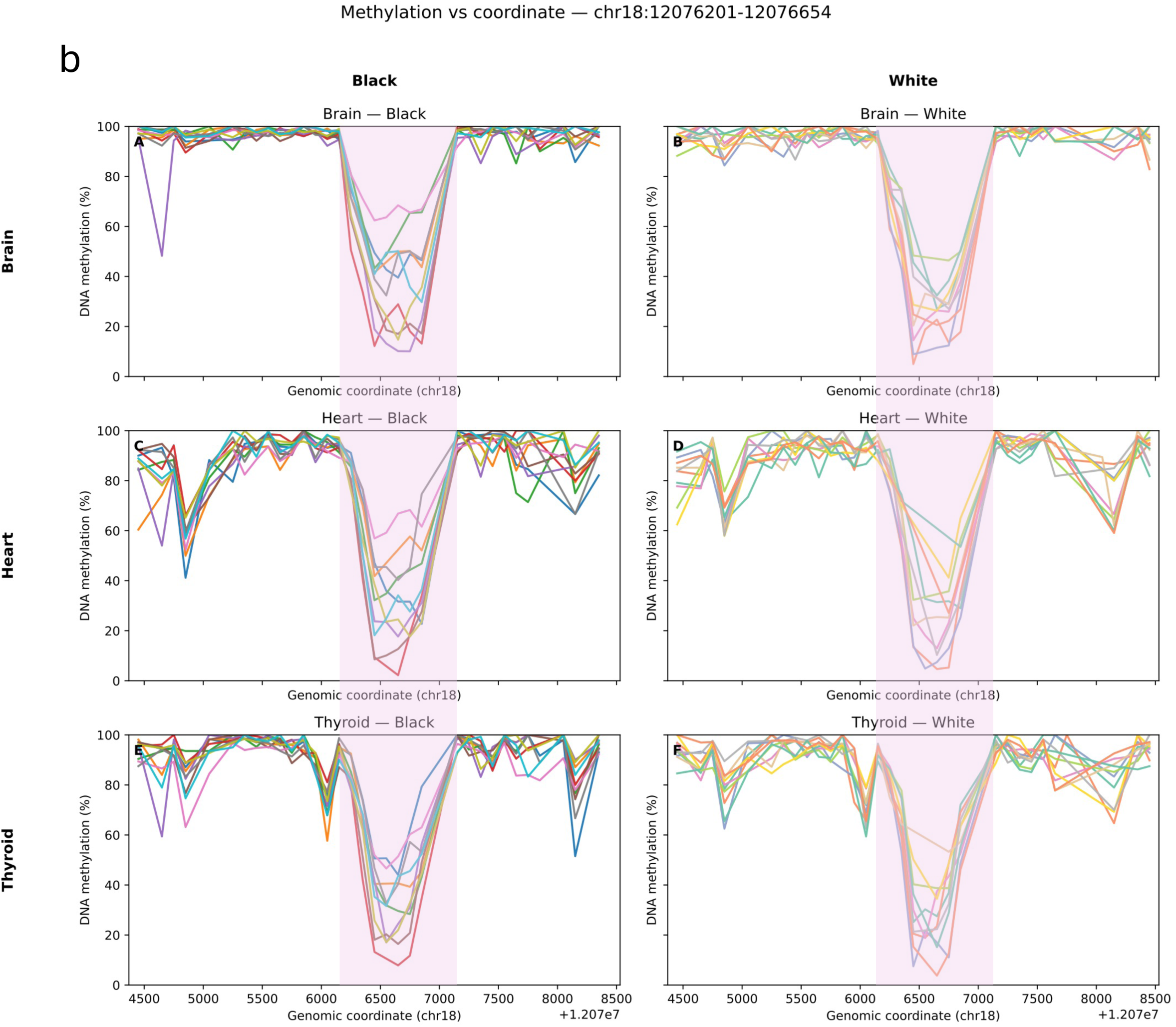
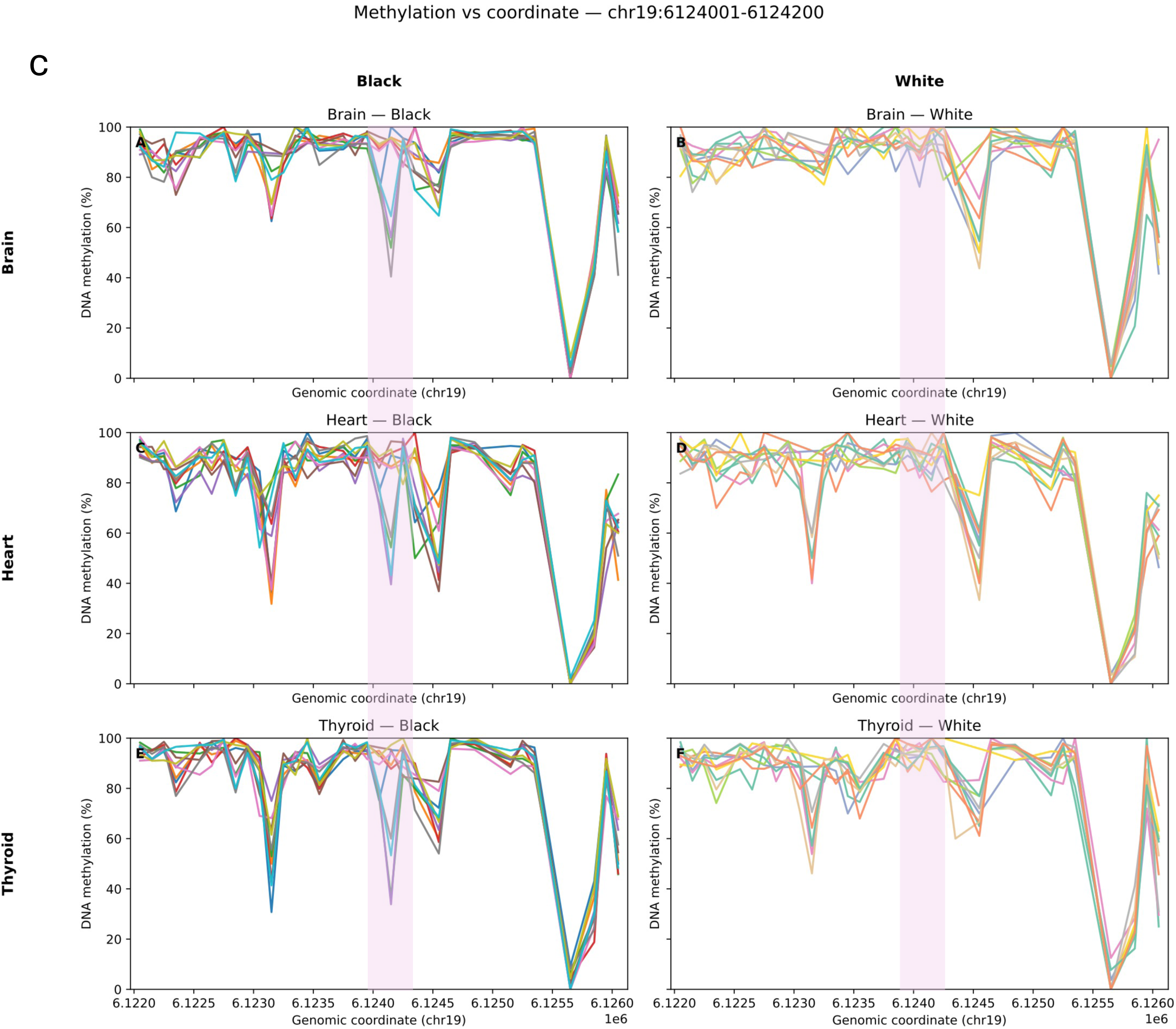
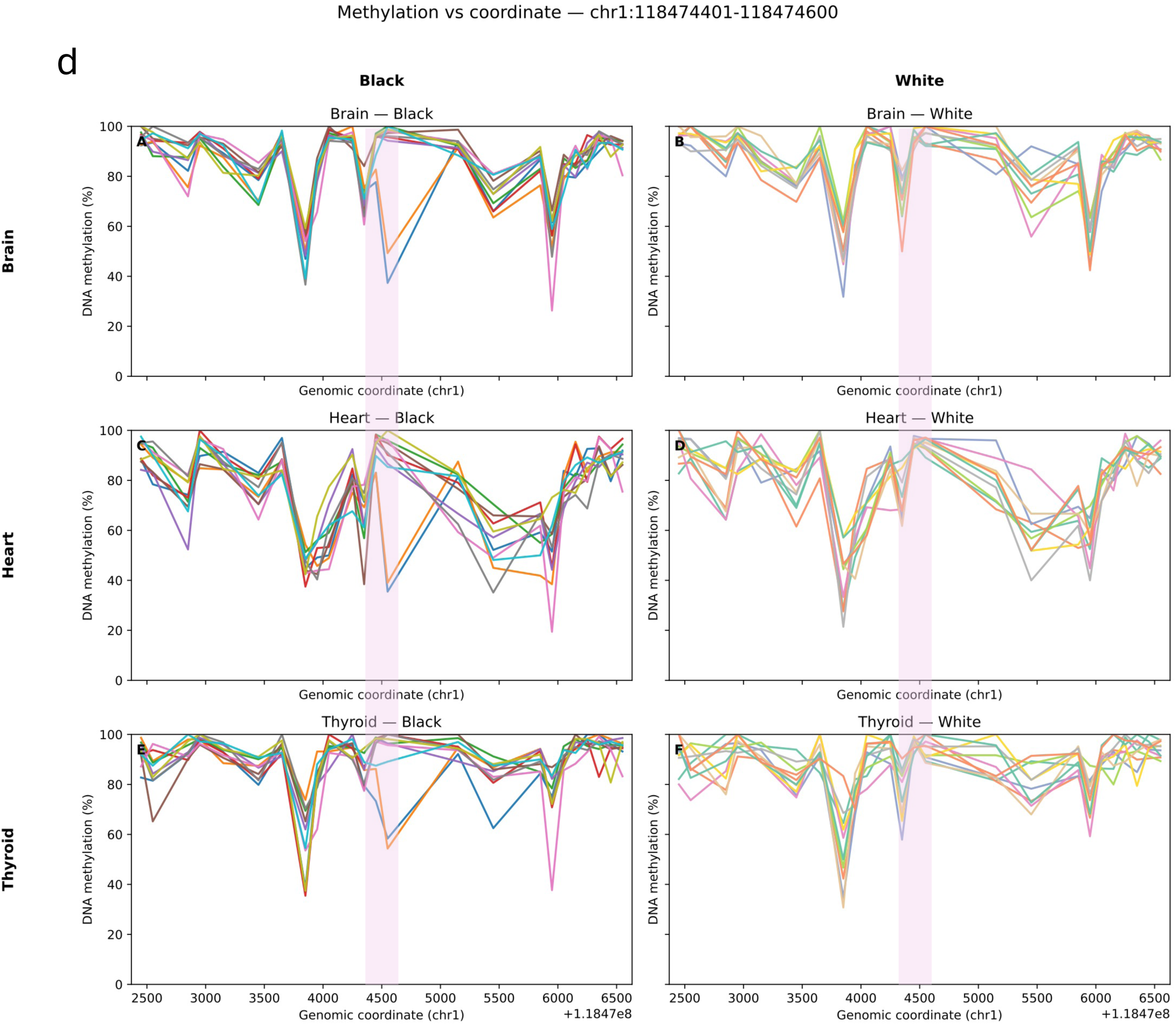
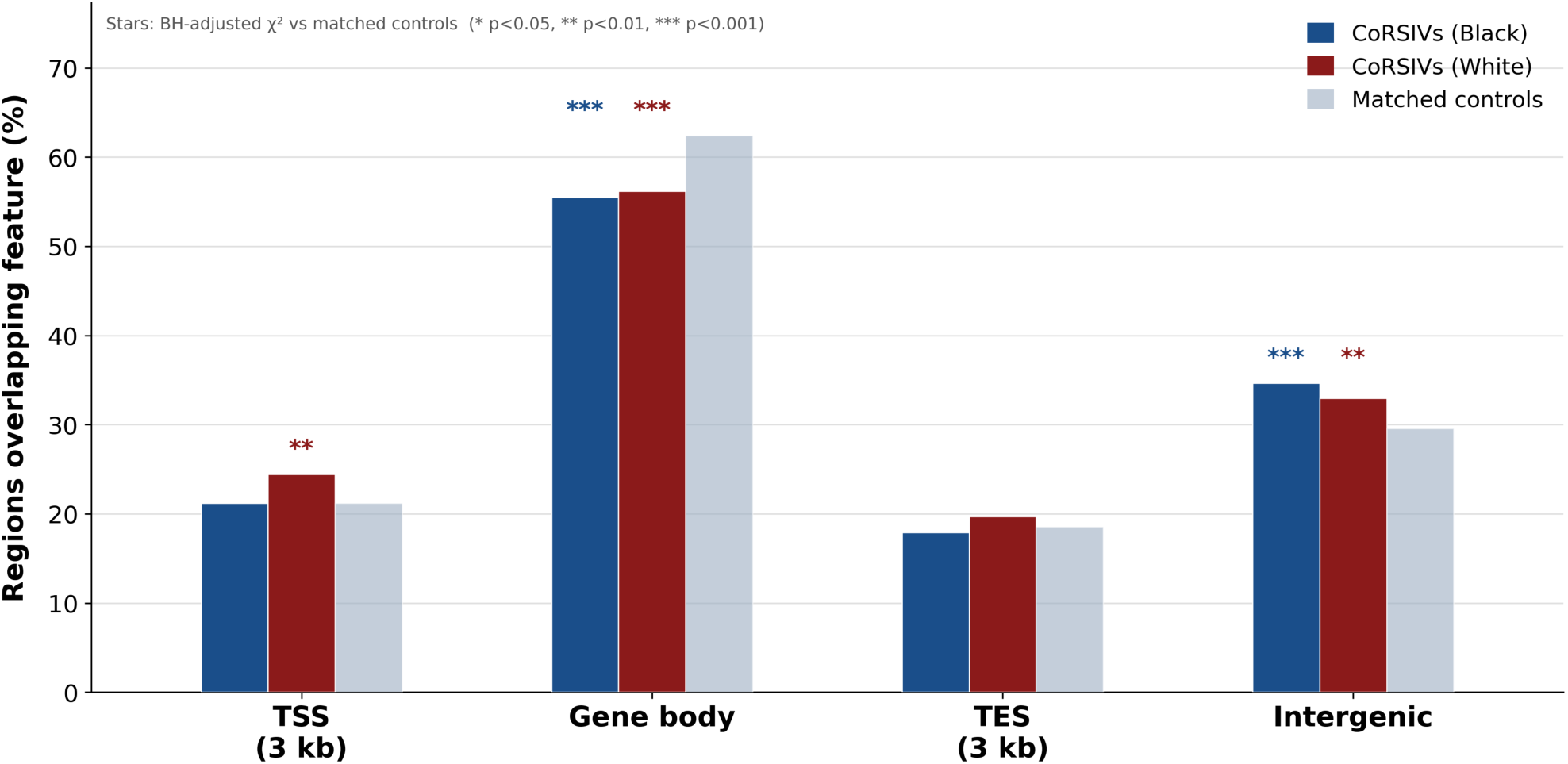
Annotation of CoRSIVs 2.0 relative to genic features. Black and White CoRSIVs, and matched control regions, were annotated using GENCODE v48 gene models and classified based on overlap with transcription start sites (TSS ±3 kb), gene bodies, transcription end sites (TES ±3 kb), or intergenic regions. Asterisks indicate statistical significance of enrichments or depletions relative to matched control regions. Although CoRSIVs are somewhat depleted in gene bodies, nearly 70% of are gene-associated.

**Figure S8.**
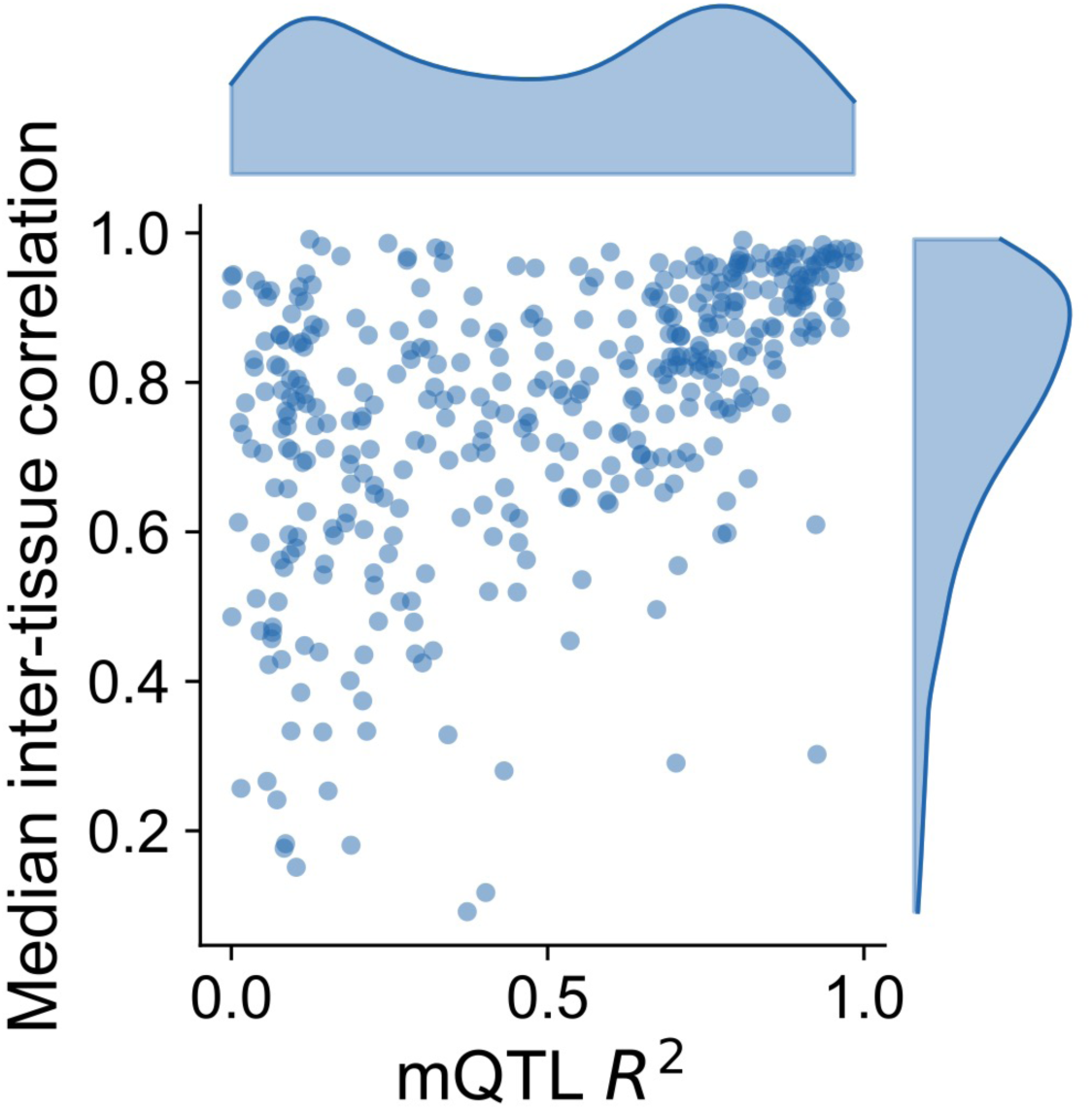
Median inter-tissue correlation vs. strength of mQTL (mQTL R2) per CoRSIV. Median ITC was calculated across all tissue pairs shown in Fig. 3d. Although CoRSIVs with strong mQTL tend to have a high ITC, there are also many CoRSIVs with high ITC despite weak mQTL.

**Figure S9.**
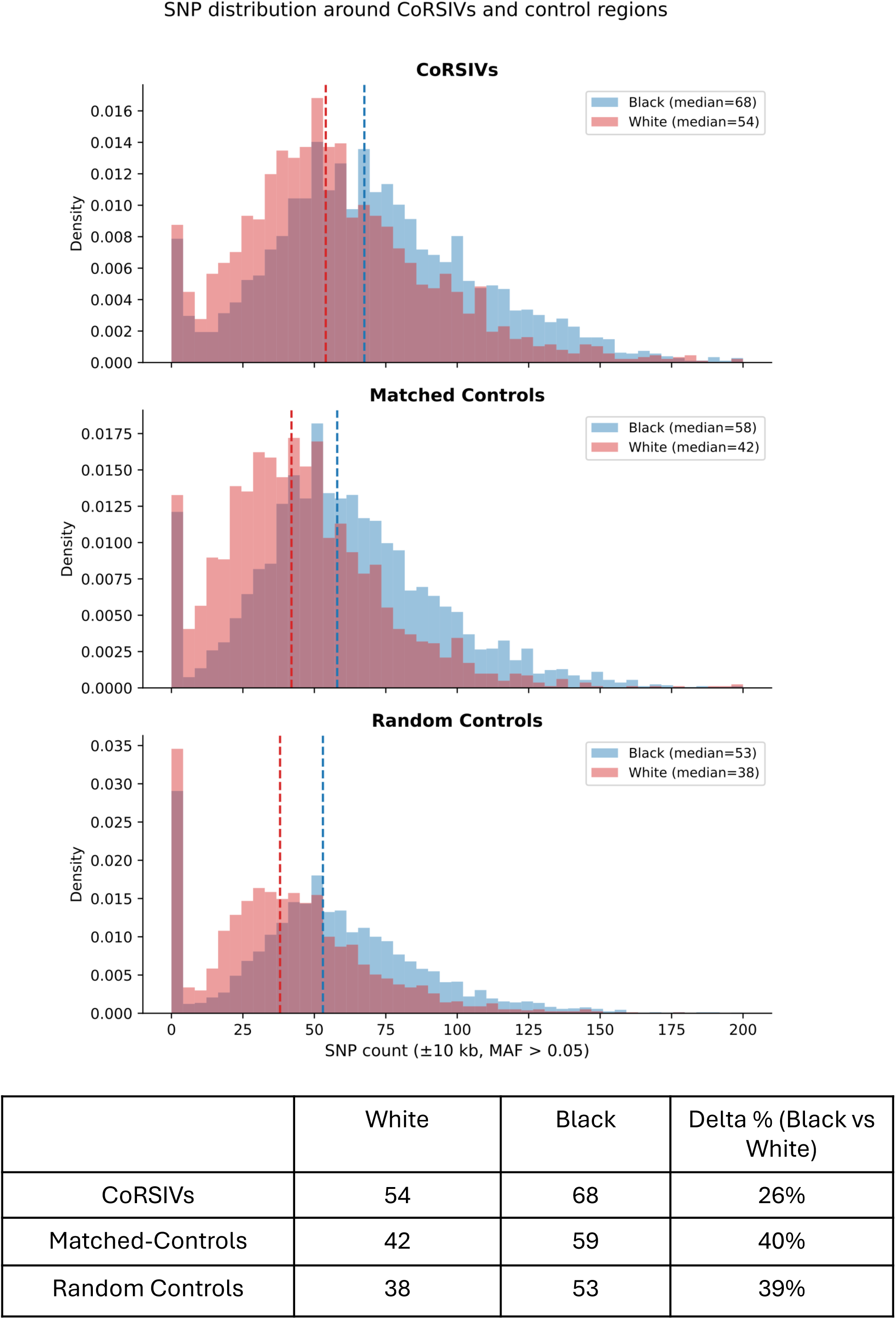
Distributions of SNV counts in the vicinity of CoRSIVs and control regions. Numbers of SNVs within ±10 kb of Black and White CoRSIVs, matched controls, and random control regions. As summarized in the table, although there are more SNVs in the vicinity of CoRSIVs relative to control regions, the excess of SNVs in the vicinity of Black vs. White CoRSIVs (26%) is less than that in control regions (∼40%).

**Figure S10.**
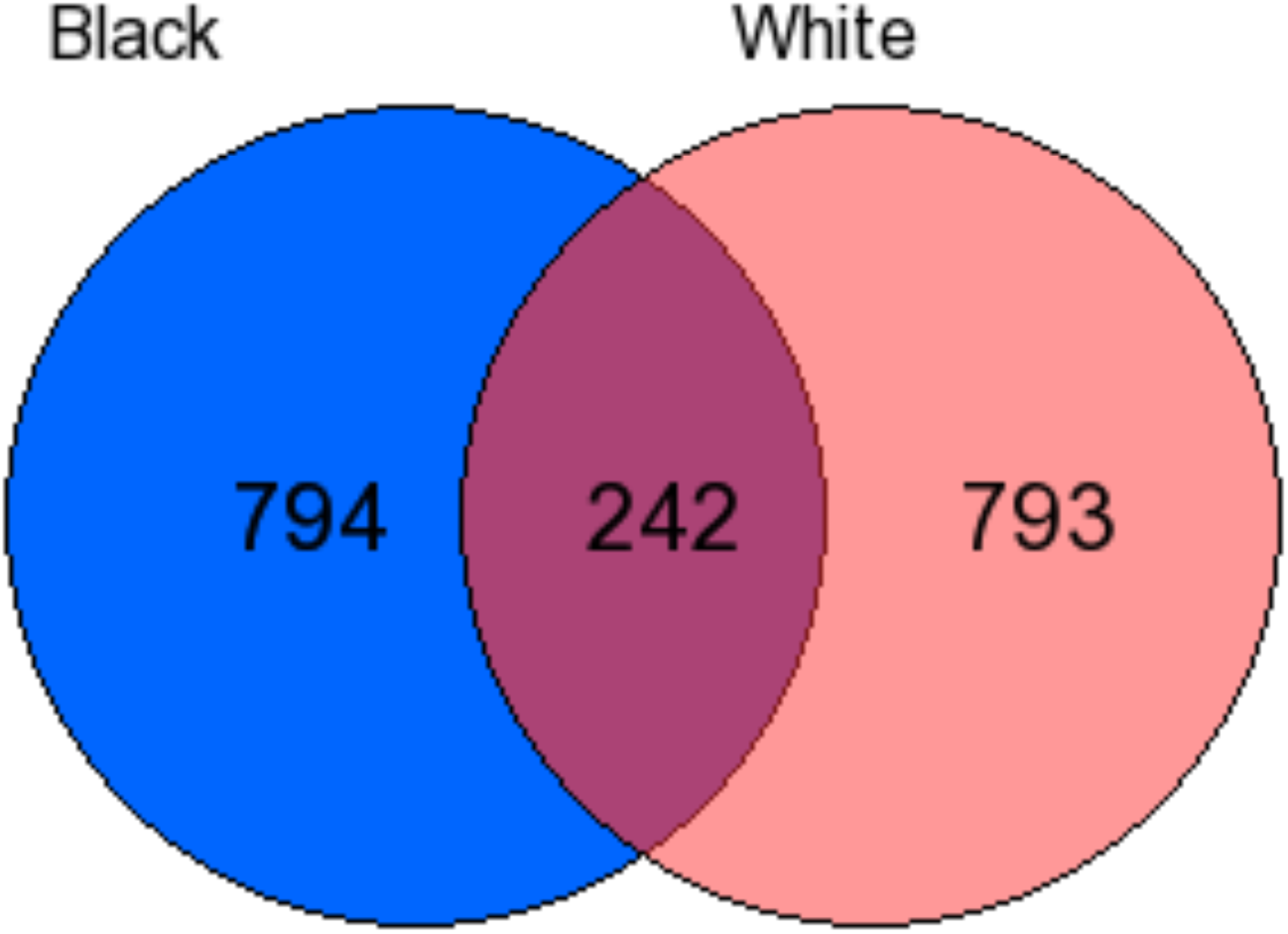
Black and White CoRSIVs largely associate with different genes. The number of genes were matched across the two sets to equalize power of the gene set enrichment analysis. Overlap of CoRSIV-associated genes is only 23%.

**Figure S11.**
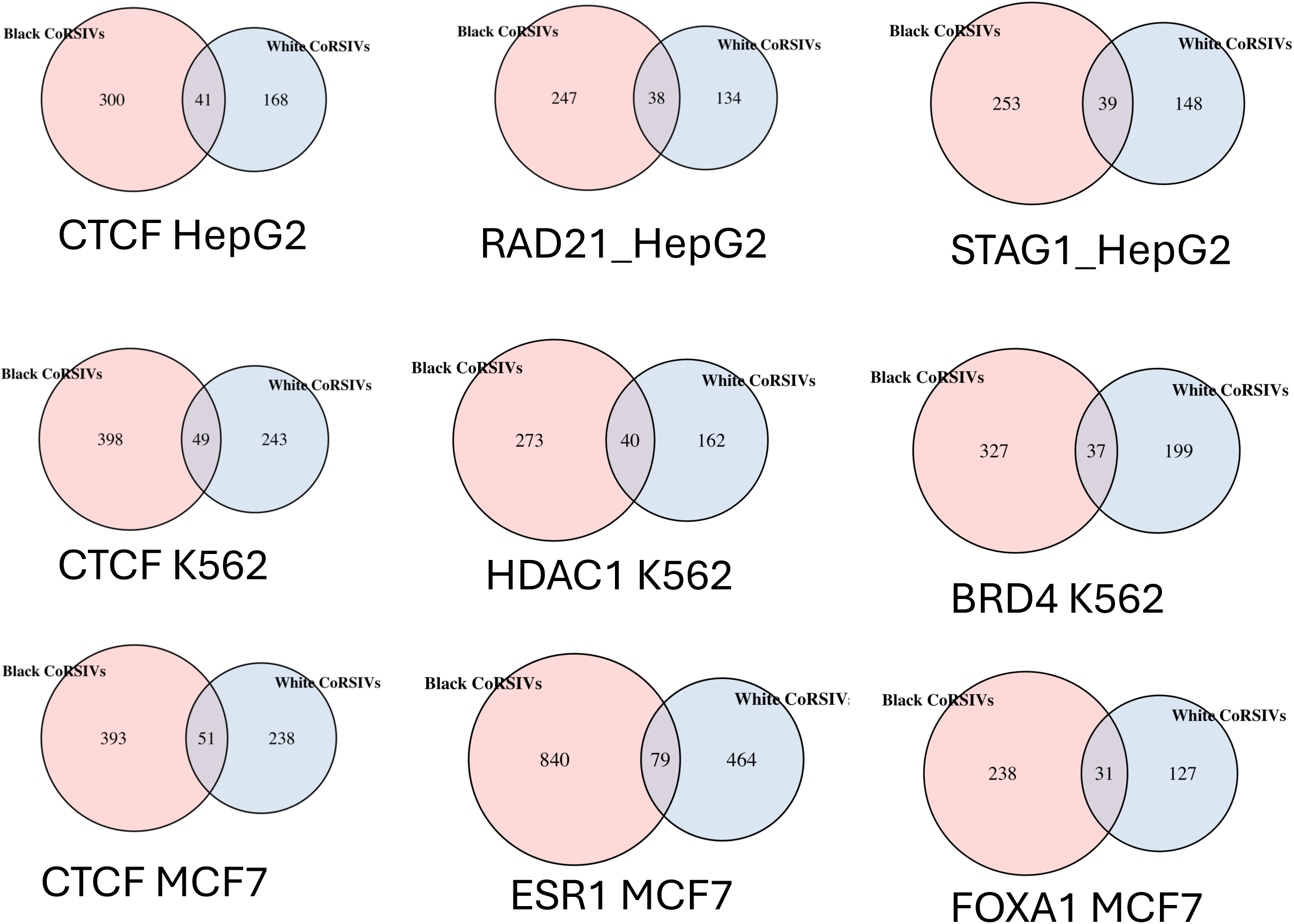
In the ReMap analysis (Fig. 5b), top TFs binding to Black and White CoRSIVs largely bind to different CoRSIVs.

**Figure S12.**
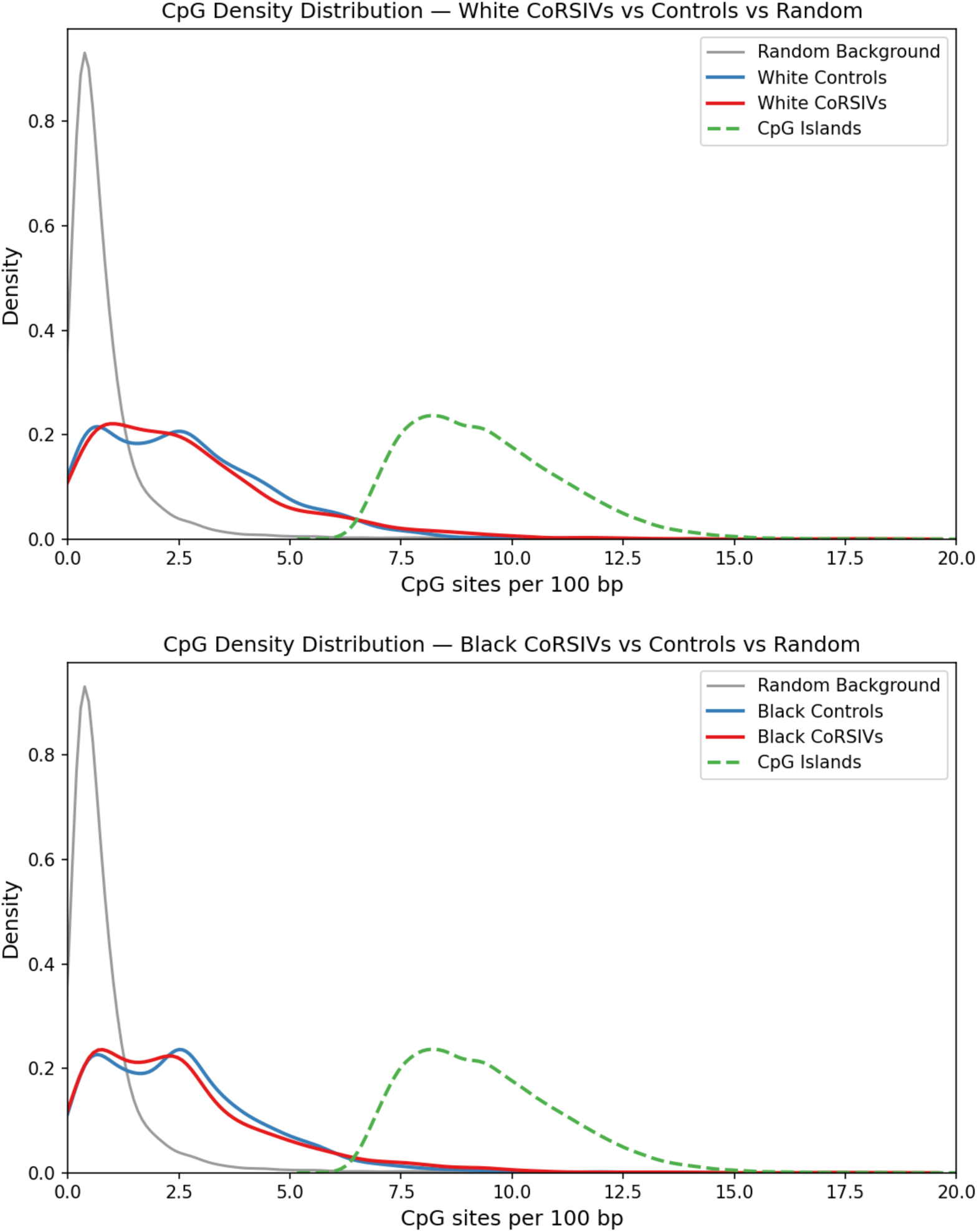
CpG density of CoRSIVs. Kernel density estimates of CpG density (CpG sites per 100 bp) are shown for White (top) and Black (bottom) CoRSIVs (red), their respective matched controls (blue), random controls (genomic background) (grey), and CpG islands (green dashed). Two thirds of CoRSIVs have a CpG density intermediate between genome background and CpG islands, an epigenetically understudied portion of the human genome.

**Figure S13.**
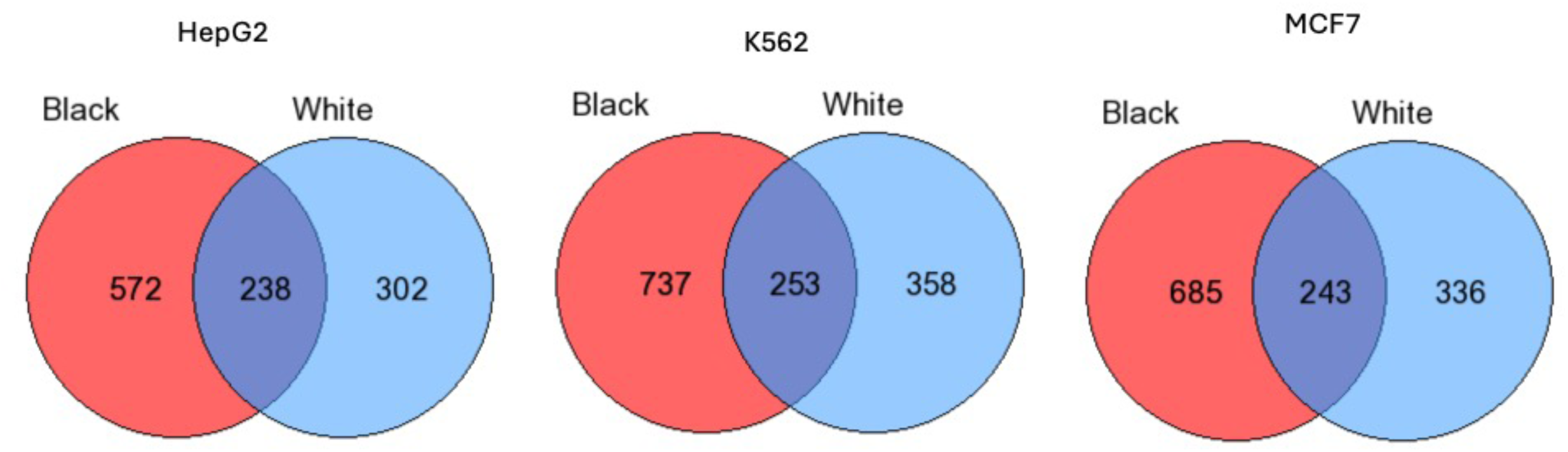
**In the gene set enrichment analysis of the ReMap data, similar terms were obtained for Black and White CoRSIVs, although the genes associated with TF binding are mostly different.**

**Figure S14.**
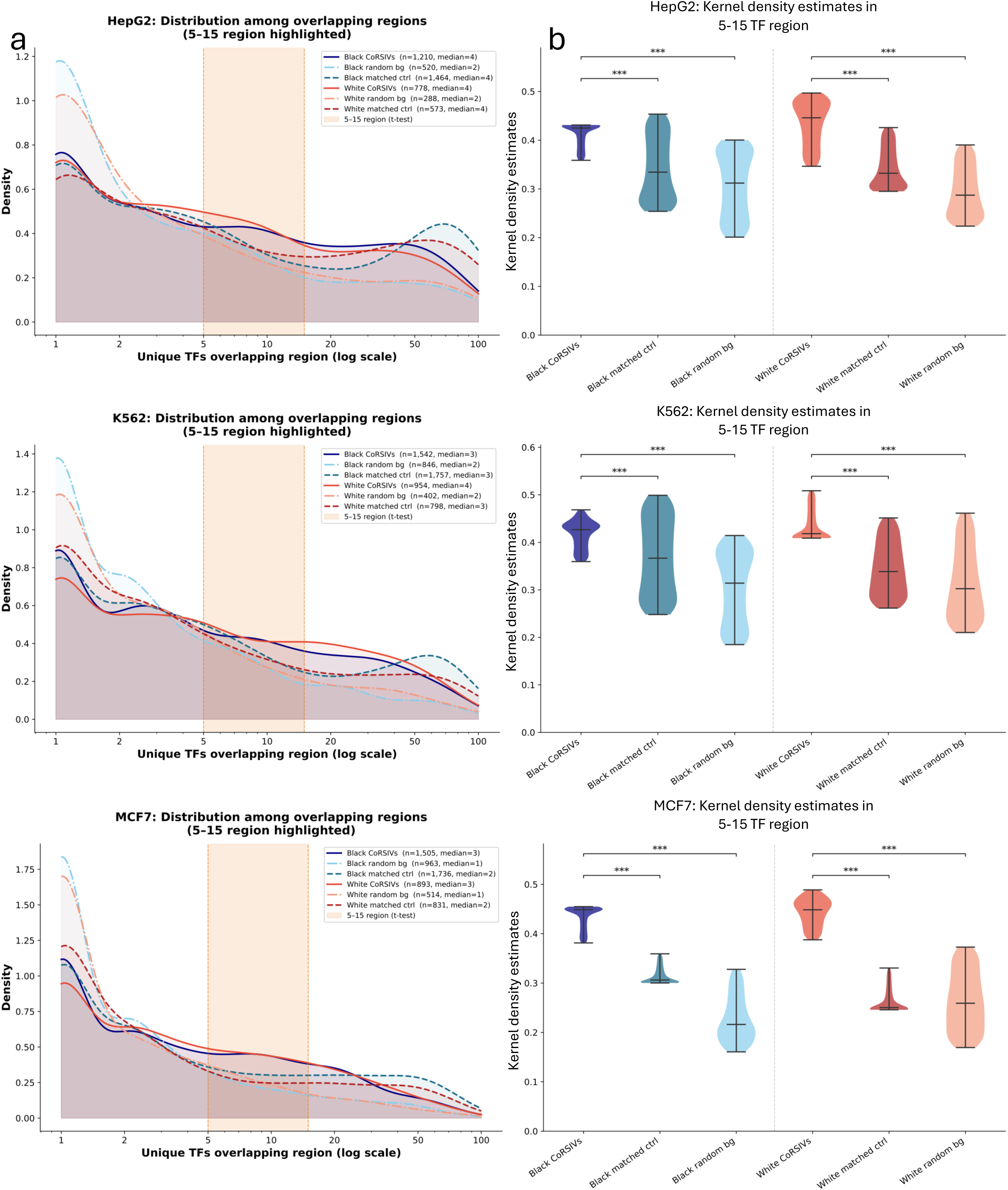
Multiplicity of TF binding per region in the ReMap data, comparing CoRSIVs, random controls, and matched controls. **a)** Density distributions summarizing the numbers of different TFs bound to each CoRSIV or control region, amongst different sets. Results are provided for the three most studied biotypes in ReMap: HepG2 (top), K562 (middle), and MCF7 (bottom). In all three, compared to random or matched control regions, each individual CoRSIV is more likely to be bound by 5-15 TFs, indicating combinatorial binding. **b)** Violin plots summarize the proportions of CoRSIVs or control regions each bound by 5-15 TFs, in the same three cell lines.

**Figure S15.**
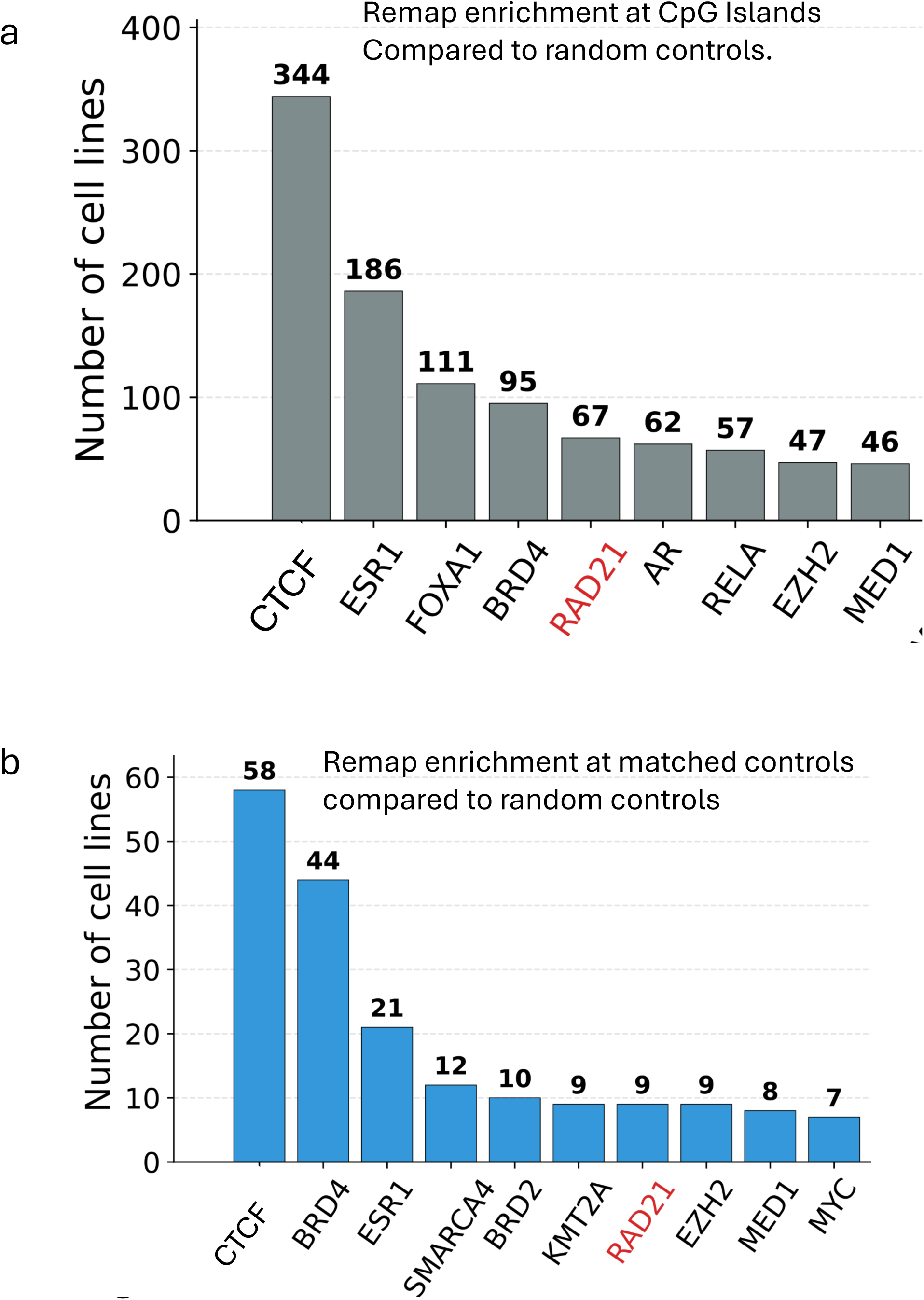
Analysis of ReMap data – top TFs binding to CpG islands and matched controls. **a)** Bar chart summarizes the top 10 TFs with evidence of enriched binding at CpG islands relative to genomic background. **b)** Bar chart summarizes the top 10 TFs with evidence of enriched binding at matched controls relative to genomic background. Similar to Fig. 5e, the most enriched TF is CTCF in both plots. But, unlike Fig. 5e, here, RAD21 is the only cohesin subunit in the top 10.

**Figure S16.**
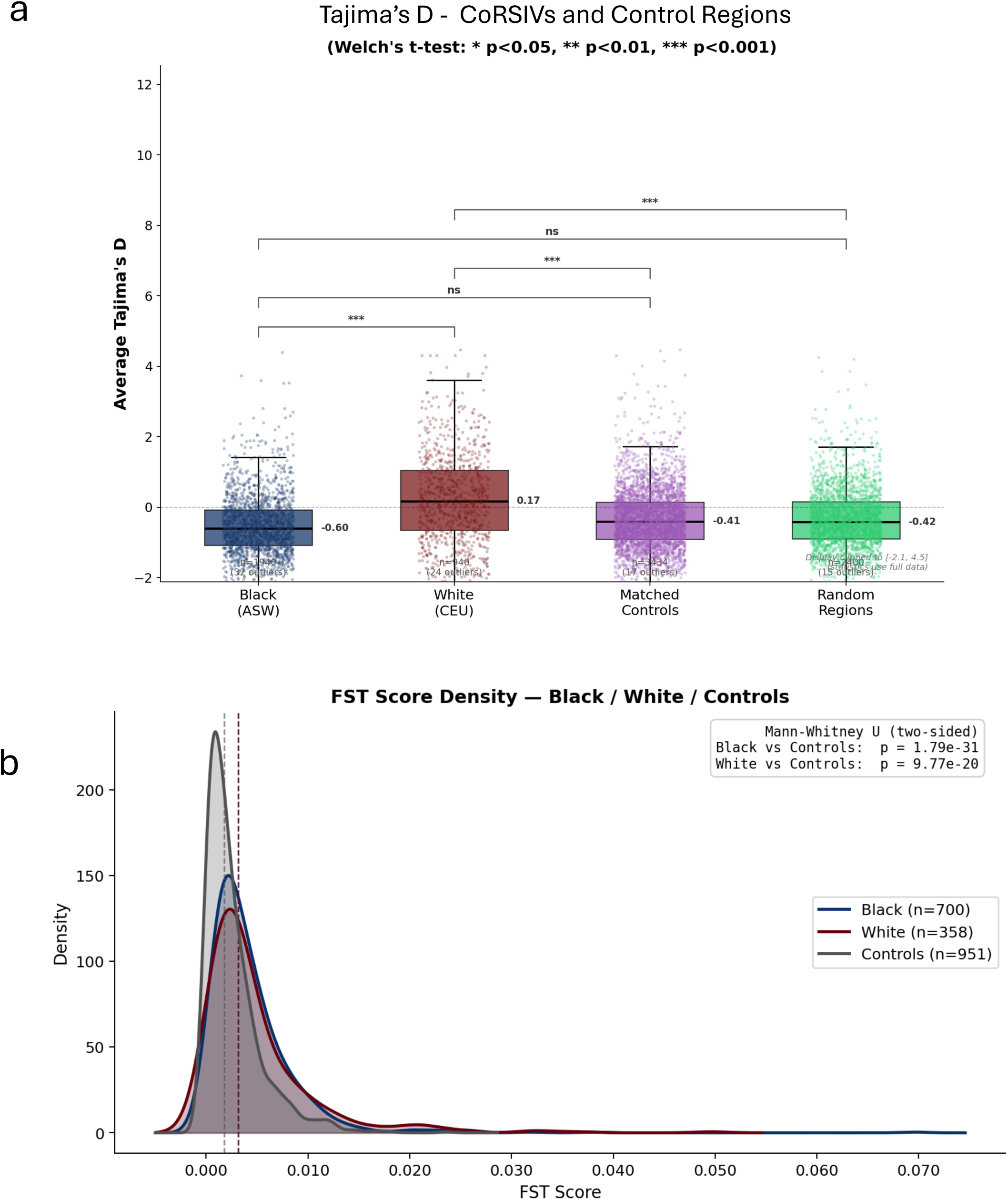
Evaluating evidence of evolutionary selection at CoRSIVs. a) Tajima’s D scores of White CoRSIVs are elevated relative to those of Black CoRSIVs, matched controls, or random control regions. b) F_ST scores are slightly higher for both Black and White CoRSIVs, relative to matched control regions.

## References

1. Gunasekara, C.J. et al. A genomic atlas of systemic interindividual epigenetic variation in humans. Genome Biol 20, 105 (2019).

2. Gunasekara, C.J. et al. Systemic interindividual epigenetic variation in humans is associated with transposable elements and under strong genetic control. Genome Biol 24, 2 (2023).

3. Dominguez-Salas, P. et al. Maternal nutrition at conception modulates DNA methylation of human metastable epialleles. Nat Commun 5, 3746 (2014).

4. Waterland, R.A. et al. Season of conception in rural gambia affects DNA methylation at putative human metastable epialleles. PLoS Genet 6, e1001252 (2010).

5. Silver, M.J. et al. Environmentally sensitive hotspots in the methylome of the early human embryo. Elife 11(2022).

6. Waterland, R.A. & Garza, C. Potential mechanisms of metabolic imprinting that lead to chronic disease. Am.J.Clin.Nutr. 69, 179–197 (1999).

7. Gunasekara, C.J. & Waterland, R.A. A new era for epigenetic epidemiology. Epigenomics 11, 1647–1649 (2019).

8. Waterland, R.A. & Michels, K.B. Epigenetic Epidemiology of the Developmental Origins Hypothesis. Annu Rev Nutr 27, 363–388 (2007).

9. Silver, M.J. et al. Independent genomewide screens identify the tumor suppressor VTRNA2-1 as a human epiallele responsive to periconceptional environment. Genome Biol 16, 118 (2015).

10. Consortium, G.T. The Genotype-Tissue Expression (GTEx) project. Nat Genet 45, 580–5 (2013).

11. Zhi, D. et al. SNPs located at CpG sites modulate genome-epigenome interaction. Epigenetics 8, 802–6 (2013).

12. Husquin, L.T. et al. Exploring the genetic basis of human population differences in DNA methylation and their causal impact on immune gene regulation. Genome Biol 19, 222 (2018).

13. Moore, J.E. et al. An expanded registry of candidate cis-regulatory elements. Nature (2026).

14. Roadmap Epigenomics, C., et al. Integrative analysis of 111 reference human epigenomes. Nature 518, 317–30 (2015).

15. Gunasekara, C.J. et al. Mouse metastable epialleles are extremely rare. Nucleic Acids Res 53(2025).

16. Derakhshan, M., Kessler, N.J., Hellenthal, G. & Silver, M.J. Metastable epialleles in humans. Trends Genet 40, 52–68 (2024).

17. Rakyan, V.K., Blewitt, M.E., Druker, R., Preis, J.I. & Whitelaw, E. Metastable epialleles in mammals. Trends Genet 18, 348–51. (2002).

18. Bryc, K., Durand, E.Y., Macpherson, J.M., Reich, D. & Mountain, J.L. The genetic ancestry of African Americans, Latinos, and European Americans across the United States. Am J Hum Genet 96, 37–53 (2015).

19. Novakovic, B. et al. Assisted reproductive technologies are associated with limited epigenetic variation at birth that largely resolves by adulthood. Nat Commun 10, 3922 (2019).

20. Peters, T.J. et al. Calling differentially methylated regions from whole genome bisulphite sequencing with DMRcate. Nucleic Acids Res 49, e109 (2021).

21. Estill, M.S. et al. Assisted reproductive technology alters deoxyribonucleic acid methylation profiles in bloodspots of newborn infants. Fertil Steril 106, 629–639 e10 (2016).

22. Boks, M.P. et al. Genetic vulnerability to DUSP22 promoter hypermethylation is involved in the relation between in utero famine exposure and schizophrenia. NPJ Schizophr 4, 16 (2018).

23. Gunasekara, C.J. et al. A machine learning case-control classifier for schizophrenia based on DNA methylation in blood. Transl Psychiatry 11, 412 (2021).

24. Chen, E.Y. et al. Enrichr: interactive and collaborative HTML5 gene list enrichment analysis tool. BMC Bioinformatics 14, 128 (2013).

25. Hammal, F., de Langen, P., Bergon, A., Lopez, F. & Ballester, B. ReMap 2022: a database of Human, Mouse, Drosophila and Arabidopsis regulatory regions from an integrative analysis of DNA-binding sequencing experiments. Nucleic Acids Res 50, D316–D325 (2022).

26. Reiter, F., Wienerroither, S. & Stark, A. Combinatorial function of transcription factors and cofactors. Curr Opin Genet Dev 43, 73–81 (2017).

27. Wray, G.A. et al. The evolution of transcriptional regulation in eukaryotes. Mol Biol Evol 20, 1377–419 (2003).

28. Ong, C.T. & Corces, V.G. CTCF: an architectural protein bridging genome topology and function. Nat Rev Genet 15, 234–46 (2014).

29. Rao, S.S. et al. A 3D map of the human genome at kilobase resolution reveals principles of chromatin looping. Cell 159, 1665–80 (2014).

30. Wang, H. et al. Widespread plasticity in CTCF occupancy linked to DNA methylation. Genome Res 22, 1680–8 (2012).

31. Meuthen, D. et al. Exploring the interplay of epigenetics and individualization. Trends Ecol Evol 41, 318–328 (2026).

32. Stajich, J.E. & Hahn, M.W. Disentangling the effects of demography and selection in human history. Mol Biol Evol 22, 63–73 (2005).

33. Bhatia, G., Patterson, N., Sankararaman, S. & Price, A.L. Estimating and interpreting FST: the impact of rare variants. Genome Res 23, 1514–21 (2013).

34. Fraser, H.B., Lam, L.L., Neumann, S.M. & Kobor, M.S. Population-specificity of human DNA methylation. Genome Biol 13, R8 (2012).

35. Heyn, H. et al. DNA methylation contributes to natural human variation. Genome Res 23, 1363–72 (2013).

36. Moen, E.L. et al. Genome-wide variation of cytosine modifications between European and African populations and the implications for complex traits. Genetics 194, 987–96 (2013).

37. Galanter, J.M. et al. Differential methylation between ethnic sub-groups reflects the effect of genetic ancestry and environmental exposures. Elife 6(2017).

38. Mozhui, K., Smith, A.K. & Tylavsky, F.A. Ancestry dependent DNA methylation and influence of maternal nutrition. PLoS One 10, e0118466 (2015).

39. Derakhshan, M. et al. Tissue- and ethnicity-independent hypervariable DNA methylation states show evidence of establishment in the early human embryo. Nucleic Acids Res 50, 6735–6752 (2022).

40. Genomes Project, C., et al. A global reference for human genetic variation. Nature 526, 68–74 (2015).

41. Feinberg, A.P. & Irizarry, R.A. Evolution in health and medicine Sackler colloquium: Stochastic epigenetic variation as a driving force of development, evolutionary adaptation, and disease. Proc Natl Acad Sci U S A 107 **Suppl 1**, 1757–64 (2010).

42. Schaub, A.M. et al. A systematic review of genome-wide analyses of methylation changes associated with assisted reproductive technologies in various tissues. Fertil Steril 121, 80–94 (2024).

43. Li, Y. et al. MAT1A Suppression by the CTBP1/HDAC1/HDAC2 Transcriptional Complex Induces Immune Escape and Reduces Ferroptosis in Hepatocellular Carcinoma. Lab Invest 103, 100180 (2023).

44. Wilson, B.G. & Roberts, C.W. SWI/SNF nucleosome remodellers and cancer. Nat Rev Cancer 11, 481–92 (2011).

45. Luijk, R., Goeman, J.J., Slagboom, E.P., Heijmans, B.T. & van Zwet, E.W. An alternative approach to multiple testing for methylation QTL mapping reduces the proportion of falsely identified CpGs. Bioinformatics 31, 340–5 (2015).

